# The mechanism of peptidoglycan O-acetylation in Gram-negative bacteria typifies bacterial MBOAT-SGNH acyltransferases

**DOI:** 10.1101/2024.09.17.613324

**Authors:** Alexander C. Anderson, Bailey J. Schultz, Eric D. Snow, Ashley S. Brott, Stefen Stangherlin, Tyler Malloch, Jalen R. London, Suzanne Walker, Anthony J. Clarke

**Affiliations:** Department of Molecular and Cellular Biology, University of Guelph, Guelph, Ontario Canada N1G 2W1; Department of Microbiology, Blavatnik Institute, Harvard Medical School, Boston, MA, USA; Department of Chemistry & Biochemistry, Wilfrid Laurier University, Waterloo, Ontario Canada N2L 3C5

**Author notes:** Anthony J. Clarke, Faculty of Science, 75 University Ave. W, Waterloo, Ontario N2L 3C5 Canada **Email:**. Michael DeGroote Institute for Infectious Disease Research, McMaster University, Hamilton, ON L8S 4K1, Canada Department of Biochemistry and Biomedical Sciences, McMaster University, Hamilton, ON L8S 4K1, Canada. Department of Chemistry, University of Waterloo, Waterloo, Ontario Canada N2L 3G1 (Stefen Stangherlin). These authors contributed equally (Bailey J. Schultz, Eric D. Snow). **Author Contributions:** A.C.A., B.J.S., E.D.S., A.S.B., S.W., and A.J.C. conceived all experiments; A.C.A., B.J.S., E.D.S., A.S.B.,S.S., and J.L. performed: gene cloning, site-directed mutagenesis and expression experiments, purification of recombinant proteins, enzyme kinetic and MS analyses, and made figures and tables; A.C.A and A.S.B. performed the X-ray crystallographic analyses; A.C.A. and T.M. performed *in silico* protein structure modelling; A.J.C. oversaw the work of A.C.A., A.S.B., S.S. and T.M.; S.W. oversaw the work of B.J.S., E.D.S., and J.L.; A.C.A., B.J.S., E.D.S., S.W., and A.J.C. wrote the paper and all reviewed the final draft.

**Keywords:** Bacterial cell wall, peptidoglycan;, O-acetylation, *O-*acetyltransferase, X-ray crystallography

## Abstract

Bacterial cell envelope polymers are commonly modified with acyl groups that provide fitness advantages. Many polymer acylation pathways involve pairs of membrane-bound *O*-acyltransferase (MBOAT) and SGNH family proteins. As an example, the MBOAT protein PatA and the SGNH protein PatB are required in Gram-negative bacteria for peptidoglycan O-acetylation. The mechanism for how MBOAT-SGNH transferases move acyl groups from acyl-CoA donors made in the cytoplasm to extracellular polymers is unclear. Using the peptidoglycan *O*-acetyltransferase proteins PatAB, we explore the mechanism of MBOAT-SGNH pairs. We find that the MBOAT protein PatA catalyzes auto-acetylation of an invariant Tyr residue in its conserved C-terminal hexapeptide motif. We also show that PatB can use a synthetic hexapeptide containing an acetylated tyrosine to donate an acetyl group to a peptidoglycan mimetic. Finally, we report the structure of PatB, finding that it has structural features that shape its activity as an *O*-acetyltransferase and distinguish it from other SGNH esterases and hydrolases. Taken together, our results support a model for peptidoglycan acylation in which a tyrosine-containing peptide at the MBOAT’s C-terminus shuttles an acyl group from the MBOAT active site to the SGNH active site, where it is transferred to peptidoglycan. This model likely applies to other systems containing MBOAT-SGNH pairs, such as those that O-acetylate alginate, cellulose, and secondary cell wall polysaccharides. The use of an acyl-tyrosine intermediate for MBOAT-SGNH acyl transfer is also shared with AT3-SGNH proteins, a second major group of acyltransferases that modify cell envelope polymers.

## Introduction

Bacteria produce a wide array of polysaccharide materials as regular components of the cell envelope^1^. Although some envelope structures, like the peptidoglycan (PG) cell wall, are highly conserved across bacterial taxa^2^, others are highly diverse even at the level of species and strain^3^. Many envelope polymers are modified enzymatically, and these modifications serve to enhance bacterial fitness through boosting antimicrobial resistance^4–6^, regulating polymer turnover and metabolism^7,8^, or limiting the host immune response to these molecules^9–11^.

The O-acetylation of PG at the C6 hydroxyl of *N*-acetylmuramyl (MurNAc) residues is a widespread modification^12^. In Gram-positive bacteria, where the cell wall is exposed to the extracellular environment, PG O-acetylation confers resistance to lysozyme and other exogenous lytic enzymes^4^ and correlates with virulence and survival in the host^13,14^. A single gene, *oatA*, is necessary and sufficient for PG O-acetylation in Gram-positive organisms^13^. The mechanism of OatA activity has been delineated, whereby an N-terminal membrane domain belonging to the acyltransferase 3 (AT3) family uses acetyl-Coenzyme A (acetyl-CoA) as a donor to auto-acetylate an invariant Tyr residue on a peptide loop at the surface of the cytoplasmic membrane^15^. The C-terminal extracellular domain of OatA, which belongs to the SGNH hydrolase family, then uses a conserved Ser-His-Asp catalytic triad to transfer this acetyl from the Tyr on the membrane domain to the Ser of the extracytoplasmic domain, and then directly onto PG. Other polymer-modifying pathways in bacteria encode fused AT3 acyltransferase and SGNH hydrolase domains, including *O*-acetyltransferases for lipooligosaccharides and lipopolysaccharides^16^.

The PG O-acetylation machinery in Gram-negative bacteria is genetically and structurally distinct from that in Gram-positive organisms^12,17^. A three-gene operon, encoding the PG *O*-acetyltransferases A (PatA) and B (PatB) and the *O*-acetyl PG esterase 1 (Ape1), is responsible for the *O*-acetyl PG phenotype^18^. Rather than belonging to the AT3 family like the N-terminal domain of OatA, PatA belongs to the evolutionarily unrelated membrane-bound *O*-acyltransferase (MBOAT) family. Both PatB and Ape1 belong to the SGNH hydrolase family. Genetic experiments have identified *patA* and *patB* as necessary and sufficient for PG O-acetylation^19^, while *ape1* mutants exhibit a hyper-*O*-acetyl PG phenotype. This is consistent with the characterization of Ape1 as a de-*O*-acetylase^17–19^. Gram-negative PG O-acetylation regulates the activity of endogenous autolysins involved in PG remodeling during growth and division^20^. Although the mechanism of PatB as a direct PG *O*-acetyltransferase has been characterized^21–23^, little is known about the mechanism of PatA beyond that the putative catalytic histidine residue conserved across all MBOAT proteins^24^ is required for its function **(Fig. 1A)**^19^.

**Figure 1.**
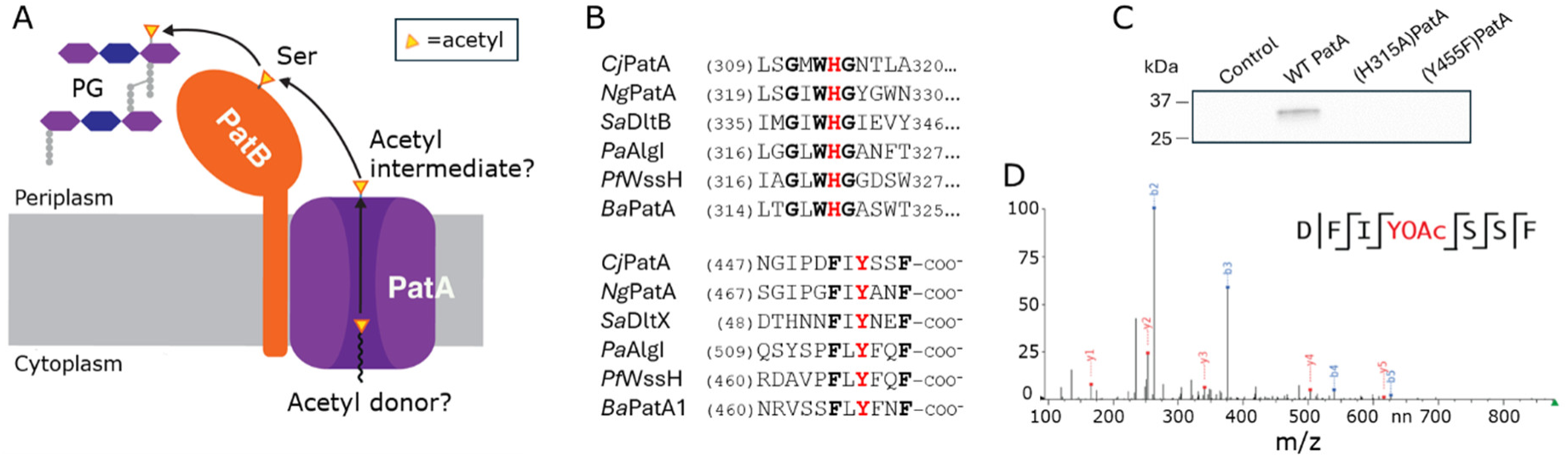
PatA forms a covalent acetyl-Tyr intermediate. (A) Previous model for PatAB-catalyzed O-acetylation of PG. It was known that PatB transfers acetyl groups to PG *via* an acetyl-Ser intermediate, but the donor of acetyl to the pathway and how PatA acetylates this Ser on PatB was unknown. (B) Partial multiple sequence alignment of bacterial MBOAT proteins encoded with SGNH proteins. Both the His and Tyr residues essential for PatA activity are strictly conserved. *Cj, C. jejuni; Sa, S. aureus; Pa, P. aeruginosa; Pf, P. fluorescens; B. anthracis.* (C) SDS-PAGE autoradiography of PatA variants incubated in the presence of [^14^C]acetyl-CoA. (D) MS/MS sequencing analysis of O-acetylated peptide from tryptic digest of PatA incubated with acetyl-CoA.

The mechanism of a related bacterial cell envelope polymer acylation system, that of the Dlt pathway for D-alanylation of (lipo)teichoic acids, was recently elucidated^25^. Like in Gram-negative PG O-acetylation, teichoic acid D-alanylation relies upon the activity of an MBOAT protein and an SGNH protein, called DltB and DltD, respectively^26^. However, instead of using acyl-CoA as a substrate, DltB uses a carrier protein containing a D-Ala thioester on its phosphopantetheinyl arm as an acyl donor. DltB transfers the D-Ala group on this carrier protein, DltC, to an invariant Tyr near the extracellular C-terminus of the membrane microprotein DltX, which binds to both DltB and DltD as part of a heterotrimeric complex. DltX then offloads the D-Ala moiety onto the catalytic Ser of DltD, which catalyzes ultimate transfer of the D-Ala residue to lipoteichoic acid. The critical Tyr of DltX is found within a highly conserved six-amino acid ‘acyl shuttle’ motif (FXYXXF). Other bacterial acylation pathways that include an MBOAT protein lack a microprotein similar to DltX. However, these MBOAT proteins contain a C-terminal motif on the final of two additional TM helices not found in DltB that is very similar to that found in DltX **(Fig. 1B)**, suggesting a shared mechanism^25^.

Here, using PatB from the human pathogens *Neisseria gonorrhoeae* and *Campylobacter jejuni* and PatA from the latter pathogen as models, we explore the mechanism of Gram-negative PG O-acetylation. We find that PatA uses acetyl-CoA to auto-acetylate the invariant Tyr residue in its conserved C-terminal motif, and we present evidence that this acetylated motif then shuttles the acyl group to PatB for transfer to PG. Importantly, MBOAT/SGNH protein pairs encoded in operons lacking *dltX*-like genes are found throughout bacteria and are responsible for acetylation of an array of cell envelope polymers. These include alginate in Gram-negative bacteria (AlgIJFX)^27,28^, secondary cell wall polysaccharides in Gram-positive bacteria (PatA1B1)^29,30^, and cellulose in both Gram-negative (WssHIFG) and Gram-positive (CcsHI) organisms^31–34^. These systems share the conserved C-terminal acyl shuttle motif found in both DltX and PatA and it is likely that they follow a similar mechanism. Although the acyltransferase domains in AT3-SGNH proteins are structurally distinct from MBOAT acyltransferases, they have also been shown to form a covalent acyl-Tyr intermediate^15^. The finding that divergent bacterial acyltransferase systems share this common mechanism suggests that transfer from a thioester to a serine ester via a tyrosyl ester is an energetically favored pathway for shuttling acyl groups to the cell envelope.

## Results

### PatA forms a covalent *O*-acetyl-Tyr intermediate

We hypothesized that PatA uses its conserved C-terminal motif to form a covalent acyl-tyrosyl intermediate^25^. To test this, we began by expressing and purifying recombinant *C. jejuni* PatA from *E. coli*. PatA is predicted to have a topology in which its N-terminus is periplasmic, so we adapted an approach used for expression of a bacterial K^+^ channel that also has an N-terminal out topology^35^. In this strategy, a tandem chimeric pectate lyase (PelB) signal sequence, maltose-binding protein (MBP), and 3C protease recognition site tag is fused to the N-terminus of PatA (**Fig. S1A**) such that the MBP is directed to fold in the periplasmic space and can be cleaved by 3C protease following purification. The construct expressed well in *E. coli,* but we could only purify low amounts of solubilized protein despite efforts to optimize the purification scheme (**Fig. S1B/C)**. Nevertheless, the amount was sufficient to test PatA activity.

To assess the activity of PatA, we incubated purified protein with [^14^C]acetyl-CoA. Enzyme reaction products were separated by SDS-PAGE and the gel was subjected to autoradiography. We observed a radioactive signal corresponding to the apparent molecular weight of PatA, implying that PatA was covalently labeled by [^14^C]acetyl-CoA (**Fig. 1C**). PatA, like other bacterial MBOATs, contains a C-terminal FXYXXF motif, and it was proposed that the invariant Tyr in this motif forms a covalent acetyl-intermediate (**Fig. 1B**).^25^ To test if PatA is acetylated at this Tyr, we generated and purified a (Tyr445Phe)PatA variant and incubated it with [^14^C]acetyl-CoA. Unlike with wildtype PatA, we did not observe radioactivity with (Tyr445Phe)PatA (**Fig. 1C**). We therefore concluded that Tyr445 in the conserved C-terminal motif is required for PatA auto-acetylation.

We also evaluated a PatA variant in which the universally conserved His residue found in all MBOAT proteins (His315 in *C. jejuni* PatA) was replaced with Ala. This equivalent His residue in DltB is required for activity^25,36^ and several other MBOAT family proteins^37–41^ (**Fig. 1B**). We did not detect a radioactive signal in the autoradiogram when (His315Ala)PatA was incubated with [^14^C]acetyl-CoA (**Fig. 1C**), showing that this His residue is required for PatA auto-acetylation. This finding is consistent with the proposed catalytic role of the invariant His, likely as a catalytic base in the transferase reaction.

To confirm the specific site of PatA auto-acetylation, we introduced unlabeled acetyl -CoA to wild-type PatA and digested the enzyme with sequencing-grade AspN aspartic protease. We analyzed the resulting peptides by LC-MS and found a distinct m/z 878.39 species corresponding to the *Cj*PatA-(452-458)-C-terminal heptapeptide sequence DFIYSSF (**Fig. S2A**). We also observed two similar peptides with an m/z of 920.40 but with increased retention times; these corresponded to the addition of a single 42.0 Da acetyl group to this same sequence (**Fig. S2B**). MS/MS fragmentation of these peptides allowed us to unambiguously assign the acetyl group in each peptide to Tyr455 and Ser456 (**Figs. 1D, S3**). Because our gel autoradiography experiment showed complete loss of radioactive signal when Tyr455 was modified, and Ser456 is not conserved in other PatB orthologs, we attribute the acetyl group on this Ser to acyl migration^42,43^ from Tyr455 during preparation of the sample for mass spectrometry. Taken together, our findings showed that PatA forms an acyl-tyrosyl intermediate in its conserved C-terminal motif.

### PatB contains a single-pass transmembrane helix

Previous work and bioinformatic predictions disagreed about whether PatB is a periplasmic or membrane protein^21^. To determine PatB’s localization, we cloned the gene encoding full-length protein from *C. jejuni* into an overexpression vector with a C-terminal His_­_tag (**Fig 2A**). Following expression in *E. coli*, we fractionated cells by osmotic shock and collected periplasmic, cytoplasmic, and membrane contents separately. Assay of these fractions for alkaline phosphatase activity indicated successful separation of our subcellular fractions (**Fig. S4**). We probed these fractions for the presence of PatB by Western blot and found it accumulated in the membrane fraction (**Fig. 2B**). We then solubilized the membrane contents with sodium lauroyl sarcosinate, a detergent selective for the inner membrane contents^44^. We observed near total recovery of PatB, indicating that PatB is an inner membrane protein (**Fig. S5A).**

**Figure 2.**
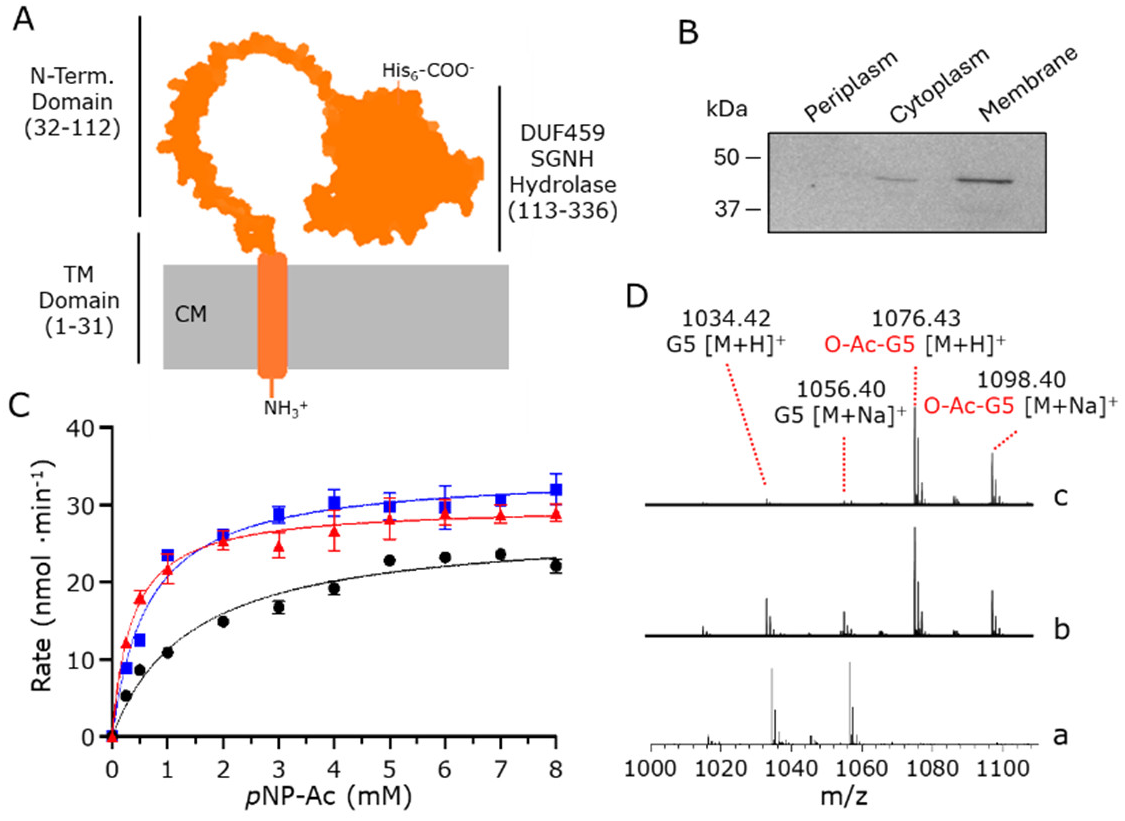
PatB is a membrane protein and its N-terminal domain is required for membrane insertion but is dispensible for activity. (A) Cartoon presentation of the AlphaFold2 model of full-length PatB toplology. Numbering in parentheses denotes residues associated with the domains. CM, cytoplasmic membrane. (B) Anti-His Western blot analysis of subcellular fractions of full length PatB from *E. coli* cells. (C) Acetylesterase activity of full-length *Cj*PatB (black), *Cj*PatB_Δ31_ (blue), and *Cj*PatB_Δ113_ (red) on *p*NP-Ac as substrate. The enzyme variants (1 µM) in 50 mM sodium phosphate pH 7.0 at 37 °C with *p*NP-Ac at the concentrations indicated. Error bars denote standard deviation (n=3). (D) LC-MS analysis of reaction products of PatB variants acting as *O*-acetyltransferases. Chitopentaose (G5) and *p*NP-Ac served as acceptor and donor substrates, respectively, for (a) no enzyme control, (b) *C*jPatB_Δ113_, and (c) *Cj*PatB_Δ31_.

We tested two N-terminally truncated variants of PatB to assess how it associates with the membrane. One variant lacked the first thirty residues (*Cj*PatB_­_), including a predicted transmembrane helix, and the other lacked the first 112 residues (*Cj*PatB_­_), including the transmembrane (TM) helix and the 82 residues preceding the SGNH domain (known as a DUF459-type SGNH hydrolase domain). Both variants accumulated as cytoplasmic proteins (**Fig. S5B**). We concluded that the first 30 residues of PatB are necessary to target it to the membrane, consistent with predictions that it contains a single-pass TM helix.

### The first 112 residues of PatB are dispensable for *O*-acetyltransferase activity

As only the 30 most N-terminal residues were necessary for PatB membrane integration, we wondered if the remainder of the 113-residue region N-terminal to the SGNH hydrolase domain might play a functional role in shaping the catalytic domain, as has been observed for the AT3-SGNH domain acyltransferase OafA involved in O-antigen acetylation^45^. We were able to purify wildtype PatB and both truncated constructs using Ni^2+^ affinity chromatography. We used our previously developed *in vitro* acetyltransferase assay^46^ to assess the kinetics of each form of *Cj*PatB. All three forms of the enzyme were active as *O-*acetylesterases and had similar kinetic parameters when the rate of acetylesterase activity was measured with *p*-nitrophenyl acetate (*p*NP-Ac; **Fig. 2C**). We also introduced chitopentaose as an acceptor into these reactions, which we have previously shown is a soluble mimetic of PG that can be used as a PatB substrate^22,23^. We observed that each form of the enzyme displayed an increased enzymatic rate relative to the donor-only reaction when chitopentaose was present at any concentration we tested, suggesting that acetyltransferase activity was present for all enzyme forms (**Fig. S6**). When we analyzed the reaction products by LC-MS, we found that we could detect an acetylated product for each enzyme form (**Fig. 2D**). Taken together, these experiments show that entire region N-terminal to the SGNH domain is dispensable for both acetyl-donor and acetyl-acceptor binding and transfer *in vitro*.

### The crystal structure of the PatB SGNH domain

To understand the mechanism of MBOAT-SGNH acyl transfer, we determined the experimental structure of PatB. Efforts to crystallize full-length *Cj*PatB or *Cj*PatB_­_were unsuccessful, but we were able to crystallize both *Cj*PatB_­_and a truncated form of *N. gonorrhoeae* PatB we previously characterized (herein *Ng*PatB_­_). We solved the initial structure of *Ng*PatB_­_by single-wavelength anomalous dispersion on a selenomethionine-labelled variant (see Methods for complete details of crystallization and structure determination) and then the structures of the native variants by molecular replacement with this structure. A complete set of X-ray diffraction and refinement statistics for our PatB structures are presented in Table S1. The structures of the SGNH catalytic domains of *Cj*PatB (PDB 8TLB) and *Ng*PatB (selenomethionine-labelled, PDB 7TLB; native, PDB 7TJB) are overall highly similar, with at most a root mean-squared deviation (RMSD) of 1.02 Å across all 217 equivalent residues between them.

The overall structure of PatB reveals a central fold that is typical of the Rossman-like SGNH hydrolase family^47^ (**Fig. 3A**). A central five-stranded parallel β-sheet (β2–4 and 7–9) is found at the core of the fold, flanked by four α-helices (α1–3 and 6). Two short, parallel β-strands (β1 and 11) fix the N- and C-terminal loops in proximity to the N-terminal side of the central β-sheet. The active site is contained in a shallow, surface-exposed depression at the C-terminal face of the central β-strand (**Fig. 3B**). A search on the DALI server for structural homologs of PatB returned a variety of known SGNH hydrolase superfamily members (**Fig. S7**).

**Figure 3.**
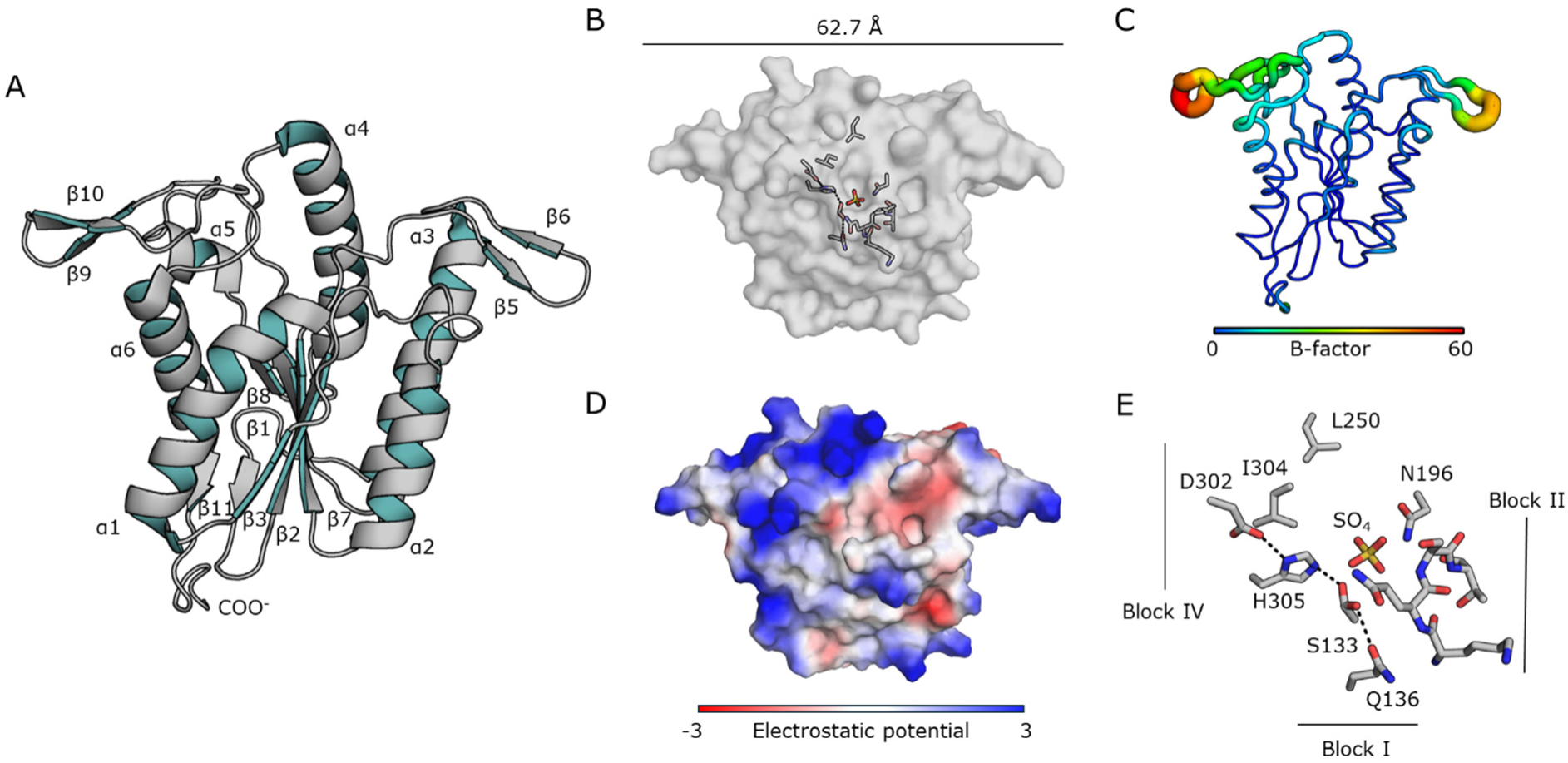
The overall structure of PatB. The native structure of *Ng*PatB_Δ100_ (7TJB) is presented as a representative of all the structural models of the PatB SGNH domain. (A) Ribbon presentation of PatB depicting a central α/β fold typical of the SGNH hydrolase family. Two extended β-hairpin motifs at the C-terminal face of the central β-sheet are present, which is not a feature of any structurally resolved SGNH hydrolase member. A short, two-stranded β-sheet joins the N- and C-terminal loops of the SGNH domain at the opposite face to the β- hairpins. (B) Surface representation showing the catalytic triad and oxyanion hole residues (as sticks) arranged on the surface of *Ng*PatB. The β-hairpin motifs extend the SGNH fold along the face that contains the active site to an approximate 60 planar surface. (C) B-factor putty model of PatB. The width and coloring of residues is based upon B-factor. The central SGNH fold contains lower (<30) B-factors, while the β-hairpin motifs display the highest B-factors (> 50). (D) The surface electrostatic potential of PatB. Several positively charged Lys and Arg residues are positioned above the active site and across the β-hairpins. (E) The active site of *Ng*PatB_Δ100_ depicting the positions of the Block I, II, and IV residues. An unusual Block II motif results in a family-atypical type I β-turn that composes the oxyanion hole, together with the Block III Asn residue. A hydrophobic “wall” is formed behind the active site. A sulfate ion is coordinated by Ser133, Ser161, Asn196 and His305.

An unusual feature of PatB is the presence of two “wing-like” β-hairpins that sit at the C-terminal face of the central β sheet that encloses the active site pocket (**Fig. 3A/B**). To our knowledge, PatB is the only structurally resolved SGNH hydrolase family member to contain these β-hairpins. These β-hairpins shape a flat, 62.7 Å wide surface on one face of PatB (**Fig. 3B**). An inspection of the crystallographic B-factors in our structures (**Fig. 3C**) demonstrated high local anisotropy about these β-hairpins, consistent with poor local electron density in our experimental maps at these locations. Interestingly, the molecular surface that they shape is rich in hydrophobic and positively charged polar residues (**Figs. 3D, S8**), and many contributing residues appear to be conserved among PatB orthologs. We suggest that these β-hairpins shape the interaction surface for PatA, PatB’s MBOAT protein partner. They may also play a role in binding the PG substrate. Interestingly, these β-hairpins are not a feature of other SGNH acyltransferases with a predicted MBOAT partner^27,29^.

The native structures of *Ng*PatB_­_and *Cj*PatB_­_contained electron density consistent with a sulfate and chloride ions in their respective oxyanion holes (**Fig. S9A,B**), while the selenomethionine variant of *Ng*PatB_­_apparently contained an empty oxyanion hole. The sulfate-bound and empty structures superimposed on each other with an all-atom RMSD of 0.12 Å, suggesting no conformational changes occurred upon ligand binding. To provide further support for this, we treated *Ng*PatB_­_with methanesulfonyl fluoride, a known covalent and irreversible inhibitor of serine proteases and esterases^48^. We observed electron density in its crystal structure (PDB 7TRR) consistent with a methylsulfonyl adduct of Ser133, the predicted catalytic residue of PatB^22^ **(Fig. S8C**). This structure superimposed on our selenomethionine-labelled, ligand-free structure also with a RMSD of 0.12 Å, suggesting no significant conformational changes occur with the formation of a covalent acyl-serine intermediate.

### The PatB active site shows a non-canonical oxyanion hole

Despite sharing low sequence identity, members of the SGNH/GDSL family are characterized by the presence of four conserved consensus motifs, referred to as Blocks I, II, III, and V^49,50^. Blocks I and V contain the catalytic triad of Ser-His-Asp, with the Ser nucleophile found within the Block I loop, while Block V loop provides the catalytic Asp and His residues (**Fig. 3E**). The previously characterized catalytic residues of *Ng*PatB_­_, Ser133, His305, and Asp302^22^ were found to be appropriately aligned to serve as the active center of the enzyme. The corresponding homologous residues in *Cj*PatB are Ser138-His310-Asp307. In both the native and selenomethionine-labelled structures of *Ng*PatB, ambiguous electron density is observed for the Oδ atom of catalytic Ser133, and so the sidechain was modelled as two alternate conformers, pointing both towards the His305 sidechain and away from it with approximately equal occupancy (0.56/0.44 and 0.55/0.45 towards/away in the selenomethionine and native structures, respectively; **Fig**. **S9A**).

The oxyanion hole is a feature of serine proteases, esterases, and lipases that is proposed to stabilize the transition states during the reaction mechanism^51,52^. In PatB, this is formed in part by the sidechain Nδ^2^ atom of Asn196 in *Ng*PatB and Asn200 in *Cj*PatB of the Block III, a conserved sequence feature of almost all families in the SGNH hydrolase superfamily^47^. However, we observed that the highly conserved Gly-containing loop residues of most SGNH hydrolase family members, which are primarily predicted or characterized esterases (i.e. non-transferases) in Block II are present as KQST in *Ng*PatB and KQNT in *Cj*PatB, with the conserved Gly replaced by Ser161 and Asn165, respectively. Structurally, this results in a type I β-turn, which also resembles the Block II found in the C-terminal domain of OatA from *S. pneumoniae*^53^, in contrast to the typical type II β-turn found in most enzymes of this family (**Fig. 3E**). The amide N atoms from this β-turn also contribute to the oxyanion hole in proximity to the Block III Asn sidechains. PatB, like OatA, shares the feature of a hydrophobic “wall” near the active site that was shown to hinder the appropriate approach of water to the active center and thereby shape activity as a transferase rather than as an esterase^53,54^. Leu250/Ile304 (*Ng*PatB) and Leu254/Val309 (*Cj*PatB) form hydrophobic patches behind the catalytic Ser from which a water molecule would need to approach the acetyl-enzyme intermediate for hydrolysis. This altered geometry and increased hydrophobicity appear to be characteristic of the known *O*-acyltransferases and may be a structural feature that delineates them from the true hydrolases/esterases in the family.

### Acetyl-tyrosine serves as an acetyl donor to PatB

We attempted to reconstitute acetyl transfer from *Cj*PatA to full-length *Cj*PatB but observed nonspecific labeling of *Cj*PatB by acetyl-CoA even in the absence of PatA (**Fig. S10**). We wondered if we could supply a minimal PatA acetyl-peptide to PatB as a substrate, or if a larger PatA surface is required to correctly align the acetyl in the PatB active site. To eliminate the possibility of acetyl group migration between the Tyr and Ser in the C-terminal motif of *Cj*PatA, we used the C-terminal peptide of *Ng*PatA with the corresponding *Ng*PatB ortholog. This ortholog contains the C-terminal sequence F473-IYAN-F478 (**Fig. 1B**), which would preclude any possibility for acetyl migration between adjacent residues. We chemically acetylated a synthetic peptide with the sequence FIYANF using acetic anhydride to yield a product with an acetyl-Tyr (**Fig. S11**). We then assessed if this peptide could serve as an acetyl donor for the O-acetylation of chitopentaose catalyzed by *Ng*PatB_­_. We found that the peptide was deacetylated in a *Ng*PatB_­_-dependent fashion (**Fig. 4A**), and that chitopentaose added to these reaction mixtures became O-acetylated (**Fig. 4B**), confirming productive acetyltransferase activity when an O*-*acetylated PatA peptide was used as a substrate for our *in vitro* assay. We also tested the minimal substrates *O*-acetyl-Ser and *O*-acetyl-Tyr as possible acetyl donors. We found that *O*-acetyl-Tyr, but not *O*-acetyl-Ser, were sufficient for transacetylation of chitopentaose by *Ng*PatB (**Fig. S12).** Along with our results that PatA can auto-acetylate its C-terminal motif Tyr (**Fig. 1C**), these experiments support our model that the C-terminal motif on PatA forms a covalent acyl-Tyr intermediate that PatB uses to acetylate PG.

**Figure 4.**
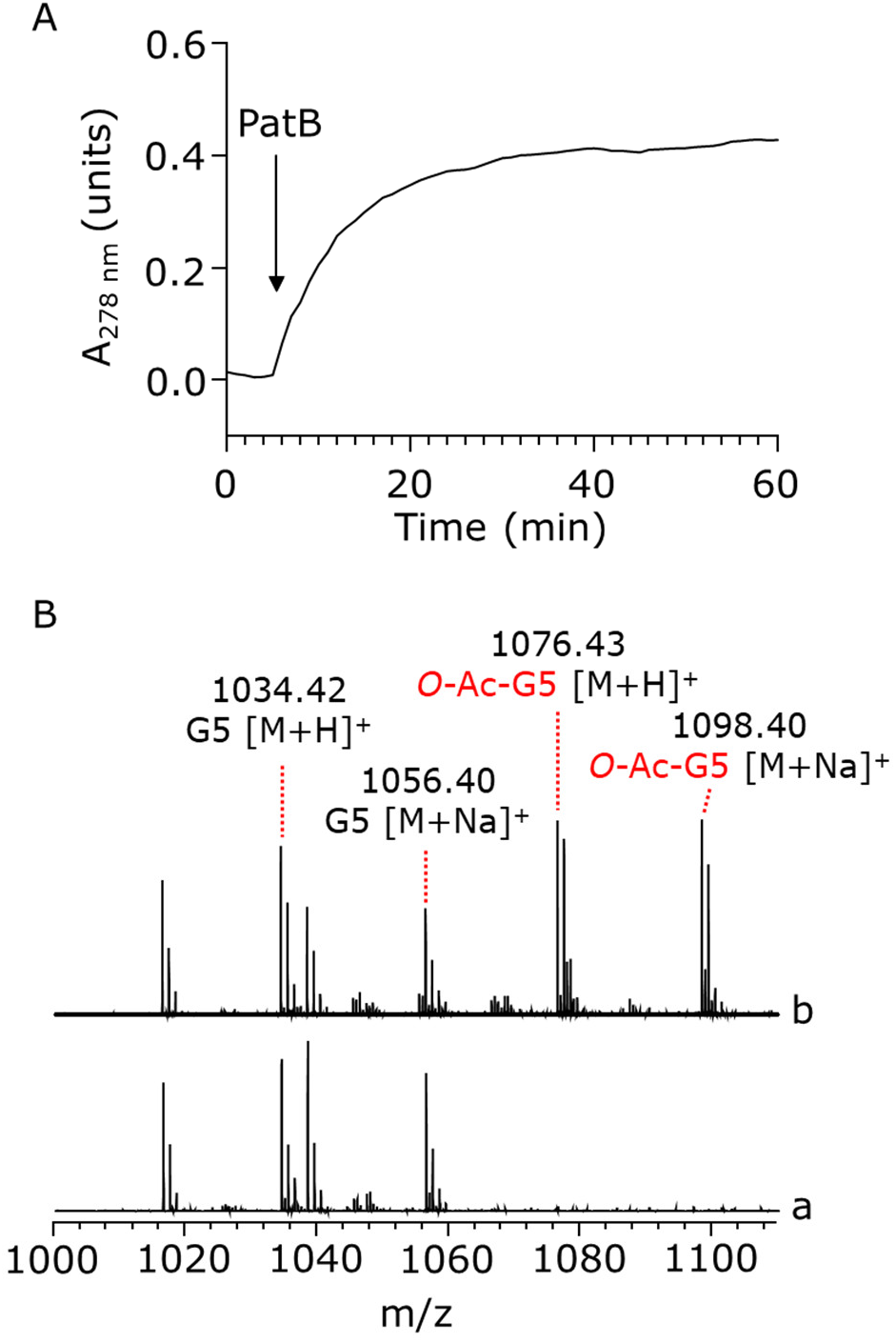
O-Acetylated C-terminal peptide of PatA as acetyl donor for PatB. (A) Esterase activity of *Ng*PatB_Δ100_ was monitored at A_278nm_ as the tyrosyl residue in the C-terminal peptide FIYANF of *Ng*PatA was de-O*-*acetylated with the addition of the enzyme (arrow). (B) LC-MS analysis of the reaction products of the transferase reaction involving incubation of *Ngj*PatB_Δ100_ with chitopentaose (G5) as acceptor in the (a) absence (negative control) and (b) presence of the O*-*acetylated peptide.

### AlphaFold Model of the PatAB Complex

Previous AlphaFold modelling of the PatAB proteins from *N. gonorrhoeae* suggested that they form a complex^25^, and we found that the same was true for the *C. jejuni* homologs. In fact, the top-scoring model predicted by AlphaFold-Multimer^55^ (**Fig. 5A**) had an overall interface template modelling score (iPTM) of 0.895 and an overall local difference distance test (pLDDT) score of 87.7 (**Fig. S13**), ranking it among the most reliable structural models of a protein complex possible with contemporary methods^55^. In the model, *Cj*PatA is an all α-helical membrane protein, typical of the MBOAT family^36^. There are 12 transmembrane passes and two re-entrant regions, totaling 17 α-helices that form a single, funnel-containing domain largely restricted to the membrane space. Helices 3-14 are part of a conserved MBOAT core that has been described for the experimentally resolved members of the family^56^. Several conserved features of MBOAT proteins are present, including a series of conserved Arg residues which line a putative acyl-coenzyme A (CoA) binding pocket at the cytoplasmic face^56,57^. A solvent-accessible lumen is formed by the core of the PatA fold that contains the putative catalytic residue His315 adjacent to the invariant Tyr455 of the C-terminal motif that sits atop the lumen near the periplasmic surface of the protein (**Fig. 5A**). The C-terminal α-helices of PatA, which do not align well with experimentally resolved MBOAT sequences, are modelled with confident scores (>70 pLDDT). The membrane-spanning helix α15 is distorted on the cytoplasmic side into a peripheral helix (α16), followed by a second transmembrane-spanning helix (α17, which carries the conserved motif linked to its terminus) back towards the periplasmic face (**Figs. 5A, S13)**.

**Figure 5.**
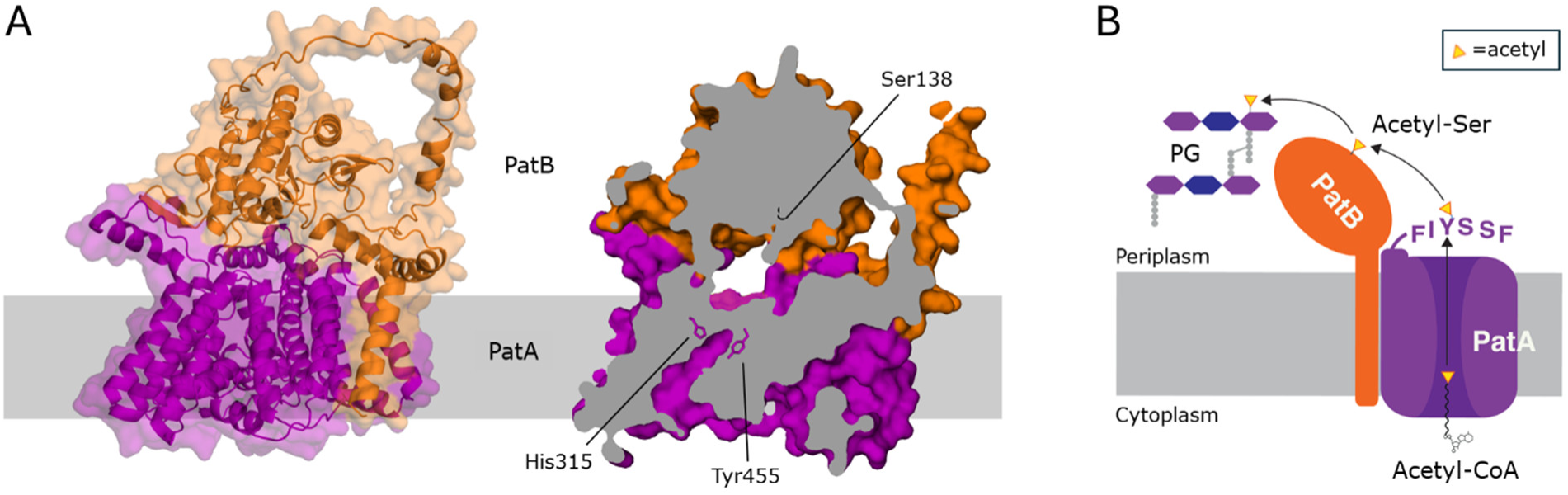
Models of the PatAB complex. (A) AlphaFold Multimer model. The overall orientation of *Cj*PatB is maintained, with the N-terminal region interacting with both the membrane and PatA (left). A cross-section (right) demonstrates that Tyr455 and His315 of PatA are positioned at the top of a solvent-accessible cavity where CoA presumably binds. Above the PatA active site is a solvent-accessible tunnel lined on top by the active site of PatB. (B) The working model for PG O-acetylation in Gram-negative bacteria. PatA transfers an acetyl group from a cytoplasmic CoA donor to a conserved periplasmic Tyr residue. PatB then transfers this acetyl group *via* a covalent Ser intermediate to PG.

The PatB N-terminal region resides in the membrane-embedded region and associates with PatA through contacts with both α1 and α17, corroborating the important role of α17 in PatA. This N-terminal region comprises three helices, with the first 32 amino acids modelled as an extended transmembrane helix with confident (> 78) pLDDT scores (**Fig. S13**). The PatB TM helix is modelled to interact with the strictly conserved Glu7 at the top of the PatA α1 via any of Tyr28, Lys32, or invariant Tyr33, modelled at distances of 2.4 to 3.4 Å. Following the TM helix are two amphipathic membrane-peripheral helices. A short coil connects the two helices, which sit on the membrane surface and are approximately perpendicular to the top of the N-terminal helix, together forming a “Y” shaped membrane-associating region. Between this Y-shaped α-helical grouping and the SGNH hydrolase domain is a region modelled as a loop with very low (< 42) pLDDT scores. The SGNH hydrolase domain itself is oriented with two β-hairpins facing toward PatA, one of which is in contact with the PatA periplasmic surface. The model of the two proteins shapes a small tunnel ranging from 10 to 18 Å wide between the two proteins, with the catalytic Ser of PatB and the catalytic Tyr/His of PatA both at opposite surfaces of this cavity (**Fig. 5A**). This positioning suggests that the C-terminal motif of PatA can move back and forth between the PatA funnel and the PatB active site, similarly to the proposed ability of the DltX C-terminal motif to move between the DltB and DltD active sites.

## Discussion

Here, we have established the mechanism for Gram-negative PG O-acetylation. Two enzymes, PatA and PatB, were known to catalyze this process^18,19^, but the chemical mechanism was unclear. We show that PatA transfers the acetyl group from acetyl-CoA to a Tyr residue in a conserved C-terminal hexapeptide motif. This motif then shuttles the acetyl group to PatB’s active-site Ser^22^, and PatB then transfers the acetyl group onto PG (**Fig. 5B**)^21–23^. Other pathways that use membrane-bound *O*-acyltransferases to acylate bacterial cell envelope polymers contain a similar C-terminal hexapeptide motif^25^ and we conclude they share the same mechanism. More broadly, it is now clear that the use of an acyl-Tyr intermediate is conserved in diverse cell envelope acylation pathways.

Acylation of extracellular polymers is a critical process in both eukaryotes and prokaryotes. In bacteria, two major systems catalyze polymer acylation. One system uses a membrane-bound *O*-acyltransferase (MBOAT) protein and a separate SGNH protein to effect acylation; the other uses a single bifunctional enzyme containing an AT3 domain fused to an SGNH domain. The teichoic acid D-alanylation (Dlt) pathway was the first mechanistically characterized acylation pathway that uses an MBOAT/SGNH pair^25^. In the Dlt pathway, D-Ala is first loaded onto a carrier protein (DltC) and is then transferred by the MBOAT protein DltB to an invariant tyrosine in the C-terminal hexapeptide of a membrane microprotein called DltX. DltX shuttles D-Ala to the SGNH protein (DltD) active site where it forms a covalent intermediate before being transferred to lipoteichoic acid^25,26^. Other MBOAT/SGNH acylation pathways lack a similar carrier protein, evidently because they use an acyl-CoA donor, as we showed here for PatA. These MBOAT/SGNH acylation pathways also lack a DltX-like membrane microprotein. However, they contain an identical C-terminal hexapeptide motif fused to the C-terminus of the MBOAT protein, suggesting they share the same chemical mechanism involving formation of an acyl-Tyr intermediate^25^. Using PatAB as a paradigm, we have now shown that the MBOAT protein PatA catalyzes auto-acetylation of the invariant Tyr residue in its C-terminal acyl shuttle motif. This acyl-Tyr intermediate serves as a donor for PatB to acetylate peptidoglycan through a covalent serine intermediate.

The first mechanistically characterized AT3-SGNH enzyme is OatA from *S. aureus*, which O-acetylates PG^4,15^. In the presence of acetyl-CoA, the AT3 domain of OatA auto-acetylates a tyrosine that then serves as the acetyl donor for the SGNH domain^15^. The SGNH domain transfers the acetyl group from this donor to PG via a covalent acetyl-Ser intermediate^53,54^. Unlike in the MBOAT/SGNH systems, where the invariant Tyr residue is found in a well-conserved motif at a protein’s C-terminus, in AT3-SGNH systems the invariant Tyr is proposed to sit between TM helices in a region of relatively low conservation. In OatA specifically, this Tyr was shown by LC-MS/MS to be acetylated when the OatA SGNH domain was inactivated by replacement of the active site Ser^15^. The evidence that AT3-SGNH proteins form an acyl-Tyr intermediate is strong and points to a conserved chemical mechanism for transmembrane acylation.

The use of a Tyr residue as a nucleophile is relatively uncommon in biology. Tyrosine phosphorylation plays a crucial role in a number of signaling cascades^58,59^ and tyrosine sulfation modulates extracellular protein-protein interactions^60^. In the context of enzymatic mechanisms, the most well-studied nucleophilic role for a Tyr residue is in DNA topoisomerases and recombinases (including Cre recombinase), where the Tyr side chain attacks phosphate in the DNA backbone to form a covalent intermediate and thereby cleave the DNA^61–64^. Additionally, nucleophilic Tyr residues are used by neuraminidases and trans-sialidases for the purpose of glycosidic bond cleavage^65–67^. In the case of the MBOAT/SGNH and AT3-SGNH fusion systems used by bacteria for cell envelope modification, we have expanded the known roles Tyr residues play to include a role as a carrier for acyl intermediates being moved across the membrane. These acyl groups begin as energetically activated thioesters in the cytoplasm generated through the use of ATP. They then move to a Tyr side chain hydroxyl group, resulting in an ester, with the help of a His residue that likely activates the Tyr for nucleophilic attack. Finally, they are transferred to their ultimate destination (an alcohol group on the polymer) by the SGNH domain, involving another covalent ester intermediate on the SGNH serine. As the acyl groups move through these systems, they move favorably from higher energy to lower energy at each step, with the pK_­_’s of the groups that house the acyl group moving from ∼9.6-9.8 (CoA/phosphopantetheine)^68^ to ∼10-10.5 (Tyr)^68^ to ∼13-14 (Ser)^69^. This thermodynamically favorable stepwise process allows for cytosolic ATP to power the movement of the acyl group without any further energy input being required. We postulate that nature has conserved this mechanism as an effective strategy to move acyl groups across the membrane and onto extracytoplasmic polymers.

The structures presented here also provide some clues to how the SGNH domains in these MBOAT/SGNH and AT3-SGNH systems maintain transferase activity over hydrolase activity. Most characterized SGNH superfamily members have activity as esterases or hydrolases^47^, but the structures of the ectodomains of OatA and now PatB suggest how this conserved fold can adopt new activities. The replacement of the Block II Gly residue, present in most SGNH hydrolases, with a Ser or Asn residue results in a type I β-turn rather than the more typical type II β-turn^47,53^. This change in secondary structure, coupled with the hydrophobic “wall” at the back of the active site in OatA and PatB, prevents the approach of a water molecule and therefore hydrolysis of the acyl-enzyme intermediate^22^. These structural features help shape the preference of these enzymes to act as transferases instead of hydrolases.

It remains unanswered whether PatAB modifies newly formed, uncrosslinked PG or mature crosslinked PG. The topology of the PatAB complex implies that PG O-acetylation must occur at the periplasmic face of the membrane. However, Lipid II, the membrane-anchored precursor for PG polymerization, is not a substrate for O-acetylation^70,71^ so the modification is installed during or after polymerization. Because PatB contains a large, presumably unstructured linker between its TM helix and soluble SGNH domain, the acetylated SGNH domain can plausibly dissociate from PatA, raising the possibility that it might modify PG strands that are crosslinked into the cell wall matrix. However, this mechanism seems unlikely because PatB does not contain any canonical carbohydrate binding domains that could target it to the cell wall. PG-binding domains are commonly found appended to autolytic enzymes with amidase, peptidase, or other PG-associated activities. For example, Ape1, the SGNH hydrolase found in the same cluster as PatAB, has a typical SGNH hydrolase fold and recognizes PG using its appended family 35 carbohydrate-binding module^72^. Assuming PatB remains associated with PatA, it seems more likely that it acts on newly formed PG near the membrane surface. AlphaFold modeling predicts that a lateral tunnel forms at the interface between PatA and PatB, and it is possible that nascent PG threads through this tunnel and is modified before being crosslinked into the sacculus.

The modification of bacterial cell envelope polymers plays important physiological roles and is often important for virulence^9–11^, colonization^10,31^, or antimicrobial resistance^4,5^. Both the MBOAT/SGNH pathways (*e.g.*, the Dlt system as well as the PatAB system studied here) and AT3-SGNH pathways (*e.g.* OatA) appear to rely upon conserved Tyr residues in the membrane component that act as nucleophiles. Furthermore we have shown that the SGNH components of both types of pathways can recognize and use peptide mimetics of the regions that contain these Tyr residues. This observation strongly implies that these pathways could be inhibited with similar peptide mimetics, which would have far-reaching consequences for bacterial physiology and may represent a viable anti-virulence strategy.

## Materials and Methods

### Bacterial strains and culture conditions

All biochemicals and reagents were purchased from either Millipore Sigma Canada (Oakville, ON) or Millipore Sigma Inc. (Burlington, MA) unless otherwise stated. The bacterial strains used in this study are listed in Table S2. Unless otherwise specified, *E. coli* was selected for and maintained at 37 °C in lysogeny broth (LB) or on agar (1.5% w/v) supplemented with carbenicillin, (100 µg/mL) ampicillin (100 µg/mL), kanamycin (50 µg/mL), or chloramphenicol (34 µg/mL) as appropriate.

### Plasmid construction

A list of plasmids and primers used in this study is presented in Table S2. PatB genes were PCR-amplified from *C. jejuni* 81-176 and *N. gonorrhoeae* FA1090 chromosomal DNA using the appropriate primers. PCR amplicons were digested with the appropriate restriction enzymes (New England Biolabs Ltd., Whitby, ON) according to the manufacturer’s protocol and ligated to the corresponding expression plasmids pre-digested with the same enzymes. Plasmids were assembled using T4 DNA ligase (NEB) according to the manufacturer’s protocol. Assembled plasmids were verified by sequencing and restriction digestion analysis. PCR-based site-directed mutagenesis of expression plasmids was performed using the protocol of Liu and Naismith^73^. Plasmids were transformed into chemically competent *E. coli* BL21(DE3) for *Ng*PatB_­_or *E. coli* C43(DE3) for *Cj*PatB derivatives using the heat-shock method^74^.

For pBS216, the entire insert of the expression construct (SS_­_through *Cj*PatA; **Fig. S1A**), plus some sequence matching pET-26b(+) on both sides (**Figs. S14, S15**), was synthesized by Twist Bioscience (South San Francisco, CA) and placed into the pTwist Amp Medium Copy vector. The insert was then amplified by PCR using primers oBS584 and oBS585, and empty pET-26b(+) was digested with NdeI and XhoI. The insert was assembled into the digested backbone using In-Fusion assembly, and the assembled plasmid was transformed into chemically competent Stellar cells. For pJL002 and pJL005, both plasmids were generated by site-directed mutagenesis of pBS216 using primers oJL003/4 (for pJL002) and oJL008/9 (for pJL005). The primers were used to amplify the entire sequence and alter the desired codon. The linear sequences were ligated using T4 polynucleotide kinase and T4 ligase and transformed into chemically competent Stellar cells.

### Protein production

For PatB expression in *E. coli*, overnight cultures were diluted 1:100 into sterile Super Broth media (32 g tryptone, 20 g yeast extract, 10 g sodium chloride per L culture). Unless specified otherwise, cultures were grown at 37 °C with aeration to an OD_­_of 0.5 to 0.8, and induction was accomplished with the addition of isopropyl β-D-1-thiogalactopyranoside (IPTG; 1 mM) for *Cj*PatB derivatives or L-arabinose (1 mM) for *Ng*PatB_­_. Induction was continued for 12–18 h at 15 °C. Following induction, cell pellets were harvested by centrifugation (5,000 x *g,* 15 min, 4 °C) and stored at -20 °C. For uniformly L-selenomethionine-labelled *Ng*PatB_­_, *E. coli* cultures were grown overnight in LB and washed thrice in M9 minimal media. The resulting culture was diluted 1:100 into M9 minimal media containing 40 mg L-selenomethionine and 40 mg all other biological L-amino acids per liter. Culture induction and harvest from minimal media were completed the same way as for rich media.

For PatA expression in *E. coli,* plasmids containing the PatA fusion construct (pBS216 and derivatives) were transformed into chemically competent C43(DE3) cells and plated at 30 °C overnight on LB-agar plates with carbenicillin. Five colonies were picked from plates and inoculated into 50 mL of Terrific Broth supplemented with carbenicillin. Overnight cultures (grown at 37 °C with aeration by shaking) were diluted 1:100 into 3 L of Terrific Broth supplemented with carbenicillin. The large cultures were grown at 37 °C with aeration by shaking. Upon reaching OD_­_values of 0.8-1.0, cultures were placed on ice for ∼10 min., expression of the PatA fusion was induced with 100 µM IPTG (final concentration), and cells were further grown with shaking at 18 °C overnight. Cells were harvested by centrifugation at 4 °C (5,000 x *g*), washed with 1x PBS, re-pelleted, flash-frozen in liquid nitrogen, and stored at -80 °C until purification.

### Protein purification

For purification of *Cj*PatA, cell pellets were thawed in an ice bath. The cell pellet (from 3 L total culture) was resuspended in 90 mL of Lysis Buffer (50 mM HEPES, pH 7.4, 500 mM NaCl, 10% (v/v) glycerol, 1 mg/mL lysozyme, 250 µg/mL Dnase I, 10 mM MgCl_­_). Resuspended cells were stirred in an Erlenmeyer flask at 4 °C for 30 min before lysis on an EmulsiFlex-C5 cell disruptor (Avestin Inc., Ottawa, ON). Cells were lysed by passaging through the EmulsiFlex-C5 at 20,000 psi for five cycles. Cellular debris were pelleted at 10,000 x *g* for 10 min at 4 °C. Membranes were isolated by ultracentrifugation of the supernatant at 140,000 x *g* for 45 min. Membrane pellets were then homogenized in homogenization buffer (50 mM HEPES pH 7.4, 500 mM NaCl, 10% (v/v) glycerol, 1% (w/v) n-dodecyl-α-D-maltoside (DDM)) using a Dounce tissue grinder (Wheaton, DWK Life Sciences, Millville, NJ). The final volume was adjusted to 40 mL with homogenization buffer, and the resulting solution was tumbled at 4 °C for 2 h. A second ultracentrifugation step at 140,000 xg for 30 min was performed to remove non-solubilized membranes. To the resulting solubilized membranes was added 1 mM imidazole (final conc.) and this was combined with 0.5 mL settled TALON resin (Takara Bio USA, Inc., San Jose, CA) that had been pre-washed with homogenization buffer. The solubilized fraction was incubated with the TALON resin for 30 min. at 4°C with end-over-end rotation. All column steps were performed at 4 °C. The batch-bound homogenate/resin mixture was added to a 25 mL disposable column, and the liquid was allowed to flow through by gravity. The resin was then washed with 10 mL Wash Buffer 1 (50 mM HEPES pH 7.4, 500 mM NaCl, 10% (v/v) glycerol, 1% (w/v) DDM, 5 mM imidazole), 10 mL Wash Buffer 2 (50 mM HEPES pH 7.4, 500 mM NaCl, 10% (v/v) glycerol, 0.2% (w/v) DDM, 10 mM imidazole), 5 mL Wash Buffer 3 (50 mM HEPES pH 7.4, 500 mM NaCl, 10% (v/v) glycerol, 0.1% (w/v) DDM, 15 mM imidazole) and 5 mL Wash Buffer 4 (50 mM HEPES pH 7.4, 500 mM NaCl, 10% (v/v) glycerol, 0.05% (w/v) DDM, 18 mM imidazole). Finally, the resin was incubated with 10 mL Elution Buffer (50 mM HEPES pH 7.4, 500 mM NaCl, 10% (v/v) glycerol, 0.05% (w/v) DDM, 150 mM imidazole), rocking at 4°C for 10 min. The resulting elution fraction was then collected. Excess imidazole was removed by repeated concentration-dilution cycles on a 50 kDa MWCO Amicon centrifugal filter unit (Amwith 50 mM HEPES pH 7.4, 500 mM NaCl, 10% (v/v) glycerol until the estimated imidazole concentration was below 2 mM.

To isolate and purify PatA from the fusion protein, in-house-purified HRV 3C protease was added to a final concentration of 1:10 (w/w) protease to fusion.Cleavage reactions were incubated overnight at 4°C with end-over-end rotation. The next day, the cleavage reactions were concentrated on a 30 kDa (MWCO) Amicon centrifugal filter unit to a volume less than 250 µL. This fraction was then further purified by size-exclusion chromatography with a pre-equilibrated Superdex 200 Increase 10/300 GL column (Cytiva, Marlborough, MA), with FPLC buffer (50 mM HEPES pH 7.4, 150 mM NaCl, 10% (v/v) glycerol, 0.05% (w/v) DDM) as the eluent. Fractions containing PatA were collected and further concentrated on a 30 kDa MWCO Amicon Ultra centrifugal filter unit down to a final concentration between 20-30 µM, as measured via Nanodrop A_­_(theoretical extinction coefficient, 8.267 x10^4^ M^-1^·cm^-1^) The final protein was aliquoted out, flash frozen in liquid nitrogen, and stored at -80 °C.

For purification of PatB proteins, *E. coli* cells were thawed and resuspended in 40 mL of lysis buffer (50 mM Tris pH 7.5, 300 mM NaCl) per L of culture, supplemented with RNase A (10 µg), DNase I (5 µg), and an EDTA-free protease inhibitor cocktail (Roche, Mississauga, ON). For purification of full-length *Cj*PatB, cells were disrupted by probe sonication at 4 °C and unbroken cells were removed by centrifugation (12,000 x *g,* 20 min, 4 °C). The membrane fraction was isolated from the lysate by ultracentrifugation (142,000 x *g,* 1 h, 4 °C). The membrane fractions were solubilized in lysis buffer supplemented with DDM (2% w/v) at 4 °C for 12–18 h with nutation. Debris were removed from the resolubilized membrane preparation by ultracentrifugation (142,000 x *g,* 1 h, 4 °C) and the resulting supernatant was loaded onto 1 mL of cOmplete Ni purification resin (Roche). The resin was washed three times in 10 mL of lysis buffer, then three times in 10 mL of wash buffer (lysis buffer plus 50 mM imidazole). The protein was eluted into elution buffer (50 mM Tris pH 7.5, 150 mM NaCl, 300 mM imidazole) and was dialyzed against 50 mM Tris pH 7.5, 150 mM NaCl at 4 °C.

For purification of *Ng*PatB_­_cells were disrupted by probe sonication at 4 °C and the insoluble fractions was removed by centrifugation (28,000 x *g,* 30 min, 4 °C). Primary purification was performed using Ni^2+^ affinity chromatography as previously described^75^, with the N-terminal His-SUMO tag removed by treatment with Ulp1 according to the protocol of Reverter and Lima^76^ and purified by ion-exchange chromatography as described previousl^75^.

For purification of *Cj*PatB_­_and *Cj*PatB_­_, cells were disrupted by probe sonication at 4 °C and unbroken cells were removed by centrifugation (12,000 x *g,* 20 min, 4 °C). The soluble fraction was isolated from the crude lysate by centrifugation (28,000 x *g,* 30 min, 4 °C). The resulting supernatant was loaded onto 2 mL of cOmplete Ni purification resin. The resin was washed three times in 10 mL of lysis buffer, then three times in 10 mL of wash buffer (lysis buffer plus 50 mM imidazole). The protein was eluted into elution buffer (50 mM Tris pH 7.5, 150 mM NaCl, 300 mM imidazole) and was dialyzed against 50 mM Tris pH 7.5, 150 mM NaCl at 4°C.

### PatA autoradiography

PatA proteins were added to a final concentration of 5 µM in a buffer containing 50 mM HEPES pH 7.4, 150 mM NaCl, and 100 µM [acetyl-1,2-^14^C]acetyl coenzyme A (100 nCi/µL; American Radiolabeled Chemicals, Inc., St. Louis, MO). Final reaction volumes were 10 µL. The reactions were incubated at 30° C for 15 min and then quenched with 3 µL of 6X SDS-PAGE loading dye. The samples were loaded onto a 4–20% polyacrylamide TGX gel (Bio-Rad) and run at 180 V until the dye front approached the bottom of the gel. Gels were removed and incubated in a solution of 40% (v/v) methanol, 5% (v/v) glycerol, and 55% water for 5–10 min. Gels were then dried in a Bio-Rad model 583 gel dryer for for 2 h and 45 min at 65° C on the slow incline setting. The dried gel was then exposed to a phosphor screen (Cytiva Multipurpose BAS-IP MS 2025 E storage phosphor screen, size 20 cm × 25 cm) for 72 h before imaging using a Typhoon FLA 9500 imager (1000 V, 50 µm).

### Trapping of the covalent PatA intermediate

PatA proteins were added to a final concentration of 5 µM in a buffer containing 50 mM HEPES pH 7.4, 150 mM NaCl, and 100 µM acetyl-CoA. Final reaction volumes were 20 µL. The reactions were incubated at 30° C for 15 min and then quenched with 6 µL of 6X SDS-PAGE loading dye. The samples were loaded onto a 4–20% polyacrylamide TGX gel (Bio-Rad, Hercules, CA) and run at 180 V until the dye front approached the bottom of the gel. The gel was removed, rinsed with MilliQ water, and stained in InstantBlue (Expedeon Ltd., Harston, UK) stain for 1 h then rinsed in water. Gel pieces with a band corresponding to PatA protein were excised, stored in MilliQ water, and sent to the Harvard Medical School Taplin Mass Spectrometry Center for modification analysis with digestion by AspN protease.

### Cell fractionation and protein detection

For cellular fractionation, the periplasmic contents of cells were isolated using the cold osmotic shock protocol of Neu and Heppel^77^. The resulting spheroplasts were disrupted by probe sonication and the membrane fraction was isolated by ultracentrifugation (142,000 x *g*, 1 h, 4 °C). Proteins were detected by SDS-PAGE and Western immunoblotting using anti-His antibodies (Invitrogen Canada Inc., Burlington, ON). Fractions were monitored for alkaline phosphatase activity by assay with 2 mM *p-*nitrophenyl phosphate. Assays (100 µL) were performed in triplicate in microtiter plates. Relative activity units were derived with the following formula:

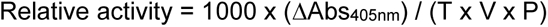

where T is time in minutes, V is assay volume in mL, and P is total protein concentration of the respective fraction (in mg/mL).

### Crystallization and structure determination

For *Ng*PatB_­_, purified protein was concentrated to 30 mg/mL. Crystallization was performed using the sitting-drop vapor diffusion method against 2.08 M ammonium sulfate and 0.1 M phosphate-citrate buffer, pH 5.5. Drops of 3 µL volume were prepared by mixing 1.5 µL each of enzyme and reservoir and equilibrated against 70 µL of reservoir. Crystals appeared after 10 to 14 days and grew to their maximum size after 30 days. Crystals were harvested and cryoprotected in 20% (v/v) ethylene glycol followed by vitrification in liquid nitrogen. For the methanesulfonyl-bound structure, 2.5 mM methanesulfonyl fluoride was added to the concentrated protein and was incubated at ambient temperature for 30 min prior to preparation of vapour diffusion experiments. Inhibition was confirmed by performing an activity assay with 2 mM *p-*nitrophenyl acetate (*p*NP-Ac) and after confirming no detectable enzymatic rate, crystallization was performed as described above. L-Selenomethionine labelled *Ng*PatB_­_crystals were grown in the same way as native crystals.

For *Cj*PatB_­_, protein was concentrated to 10 mg/mL following purification. Crystallization was performed using the hanging-drop vapor diffusion method against 100 mM HEPES:NaOH pH 7.5, 200 mM NaCl, and 25% (w/v) poly(ethylene glycol) 3500. Drops of 2 µL volume were prepared by mixing 1 µL each of enzyme and reservoir and equilibrated against 500 µL of reservoir. Crystals appeared after 48 to 72 h following drop setting and grew to their maximum size in 12 to 14 days. Harvested crystals were cryoprotected in a solution of 50% (v/v) glycerol and vitrified in liquid nitrogen. Data collection was performed on beamline 08-ID1 at the Canadian Light Source synchrotron using incident radiation with a wavelength of 0.97955 Å for native crystals and 0.97800 Å for the L-selenomethionine labelled *Ng*PatB_­_crystal. Collected data were processed with XDS^78^.

For *Ng*PatB_­_crystals, a P 3_­_space group was determined with one copy of *Ng*PatB_­_in the asymmetric unit for all crystal forms. Phases for the anomalous dataset were calculated using the AutoSol Experimental Phasing function within PHENIX^79^ using the single-wavelength anomalous dispersion method. A protein model was generated for the resulting electron density map using PHENIX AutoBuild. This model was then used to calculate the phases of the native and methanesulfonyl-bound crystal data sets via molecular replacement using Phaser-MR coupled with AutoBuild to generate a protein model for the native data set, both within the PHENIX software suite. Further refinement of both datasets was accomplished by iterative cycles of manual building/remodeling and refinements using Coot^80^ and PHENIX.Refine respectively. The refinement progress was monitored by the reduction and convergence of R*_­_*as calculated by PHENIX.

For *Cj*PatB_­_, a P 6_­_space group was determined with a single copy of *Cj*PatB*_­_*in the asymmetric unit using POINTLESS, then scaled using SCALA and data reduction performed using CTRUNCATE^71^ using a maximum resolution of 1.6 Å to ensure data completeness. The processed data were solved by molecular replacement with *Ng*PatB_­_(PDB, 7TJB) as the search model using the Phaser tool in PHENIX. Structural refinement was also performed using a maximum resolution of 1.6 Å using iterative rounds of the automated Refine tool in PHENIX, followed by manual refinement and solvent placement using Coot. Figures were prepared with PyMOL (v2.5.4).

### *In silico* modelling of PatAB

Prediction of PatB topology was performed with SignalP^81^ (v6.0) and TMHMM^82^ (v1.0.24). Modelling of PatA and PatB were performed with AlphaFold Multimer^55^ and figures were prepared with PyMOL (v2.5.4).

### Measurement of PatB rates

The assay for PG *O*-acetyltransferase was used to measure the activity of recombinant PatB proteins as described previously^46^. Briefly, *p*NP-Ac or acetyl-amino acids (1 mM) were added to PatB proteins (1 µM) at the specified concentration (acetyl donor) with or without the addition of chitopentaose (acetyl acceptor; Megazyme, Wicklow, Ireland) in 50 mM sodium phosphate pH 7.0. Measurement of chitopentaose kinetics was performed at a fixed *p*NP-Ac concentration of 2 mM, which exceeds the measured *K_­_*of all enzyme forms for this acetyl donor. Reactions (100 µL) were performed in triplicate at ambient temperature and the absorbance at 405 nm was monitored in an imaging microplate reader (BioTek, Winooski, VT) for 30 min. Enzyme-free controls were used to measure the background rates of *p*NP-Ac hydrolysis and all rates were corrected for a matched enzyme-free control. All rates reported represent the mean and standard deviation of three replicate reaction measurements. Model-fitting and kinetic parameters were derived with GraphPad Prism (v9.5.1: GraphPad Software, Boston, MA). Detection of acetylation was carried out by direct analysis of enzymatic reactions by LC-MS on an Agilent 1200 instrument coupled to an Agilent UHD 6530 Q-TOF mass spectrometer (Agilent Technologies Canada, Mississauga, ON) as described previously^29^.

### PatA C-terminal peptide preparation and analysis

The peptide H_­_N^+^-FIYANF-COO^-^ was synthesized and provided as a lyophilized trifluoroacetate salt by GenScript (Piscataway, NJ). The peptide (2–4 mM) was resuspended in 50 mM phosphate, pH 7.4 and 50% DMSO and chemically acetylated with the addition of 30 equivalents of acetic anhydride. The reaction was monitored at 278 nm until complete, typically 15 min. The resulting reaction mixture was purified directly by HPLC using an Agilent 1200 liquid chromatograph equipped with a semi-preparative C_­_column (Gemini C_­_, 10 x 250 mm, 5 µm: Phenomenex, Torrance, CA) with the following solvents: water with 0.1% formic acid (A) and acetonitrile with formic acid (B). The column was equilibrated in 2% B and elution was achieved with a linear gradient to 65% B over 35 min with monitoring at 278 nm.

Fractions were directly assessed by LC-MS and the product was identified using available MS/MS for assignment. The purified acetyl-peptide was dried under vacuum and resuspended at a final concentration of 100 µM in 50 mM phosphate buffer, pH 6.5 with 50% v/v DMSO and 1 mM chitopentaose. The reaction was monitored at 278 nm in a UV-transparent quartz glass cuvette for 5 min, and then *Ng*PatB_­_was added to a final concentration of 10 µM. The reaction was monitored at 278 nm for an additional 55 min and the product was then filtered through a 0.45 µm filter and used directly for LC-MS analysis.

## Acknowledgments

We thank Drs. Dyanne Brewer and Armen Charchoglyan of the Mass Spectrometry Facility (Advanced Analysis Centre, University of Guelph) for expert technical assistance and advice. We also thank Ross Tomaino of the Taplin Mass Spectrometry Facility at Harvard Medical School for PatA modification analysis. These studies were supported in part by operating grants from the Canadian Institutes of Health Research (TGC 114045) and the Canadian Glycomics Network (https://canadianglycomics.ca/) to A.J.C., and the National Institutes of Health to S.W. (R01 AI148752, P01 AI083214, and U19 AI158028). A.C.A. and A.S.B. were supported in part by graduate scholarships from the Natural Sciences and Engineering Research Council of Canada and the Province of Ontario (Ontario Graduate Scholarship), respectively. B.J.S. was supported in part by a graduate fellowship from the National Science Foundation (DGE 2140743), and J.R.L. was supported in part by the Harvard Summer Honors Undergraduate Research Program (NIH grant R25 HL121029) and the Harvard Division of Medical Sciences. Any opinions, findings, and conclusions or recommendations expressed in this material are those of the author(s) and do not necessarily reflect the views of the National Science Foundation. Research described in this paper was performed using the CMCF beamlines at the Canadian Light Source, a national research facility of the University of Saskatchewan, which is supported by the Canada Foundation for Innovation (CFI), the Natural Sciences and Engineering Research Council (NSERC), the National Research Council (NRC), the Canadian Institutes of Health Research (CIHR), the Government of Saskatchewan, and the University of Saskatchewan.

## References

1. Silhavy, T. J., Kahne, D. & Walker, S. The Bacterial Cell Envelope. Cold Spring Harb Perspect Biol 2, 1–16 (2010).

2. Vollmer, W., Blanot, D. & De Pedro, M. A. Peptidoglycan structure and architecture. FEMS Microbiol. Rev. 32, 149–167 (2008).

3. Whitfield, C. & Trent, M. . S. Biosynthesis and export of bacterial lipopolysaccharides. Annu. Rev. Biochem. 83, 99–128 (2014).

4. Bera, A., Herbert, S., Jakob, A., Vollmer, W. & Götz, F. Why are pathogenic staphylococci so lysozyme resistant? The peptidoglycan *O*-acetyltransferase OatA is the major determinant for lysozyme resistance of *Staphylococcus aureus*. Mol. Microbiol. 55, 778–787 (2005).

5. Kristian, S. A. et al. D-Alanylation of teichoic acids promotes group A *Streptococcus* antimicrobial peptide resistance, neutrophil survival, and epithelial cell invasion. J. Bacteriol. 187, 6719–6725 (2005).

6. Salazar, J., Alarcón, M., Huerta, J., Navarro, B. & Aguayo, D. Phosphoethanolamine addition to the heptose I of the lipopolysaccharide modifies the inner core structure and has an impact on the binding of Polymyxin B to the *Escherichia coli* outer membrane. Arch. Biochem. Biophys. 620, 28– 34 (2017).

7. Williams, A. H., Wheeler, R., Deghmane, A. E., Santecchia, I., Schaub, R. E., Hicham, S., Moya Nilges, M., Malosse, C., Chamot-Rooke, J., Haouz, A., Dillard, J. P., Robins, W. P., Taha, M. K. & Gomperts Boneca, I. Defective lytic transglycosylase disrupts cell morphogenesis by hindering cell wall de-O-acetylation in *N. meningitidis*. Elife 9, 1–23 (2020).

8. Blackburn, N. T. & Clarke, A. J. Characterization of soluble and membrane-bound Family 3 lytic transglycosylases from *Pseudomonas aeruginosa*. Biochemistry 41, 1001–1013 (2002).

9. Whitfield, G. B., Marmont, L. S. & Howell, P. L. Enzymatic modifications of exopolysaccharides enhance bacterial persistence. Front. Microbiol. 6, 1–21 (2015).

10. Pier, G. B., Coleman, F., Grout, M., Franklin, M. & Ohman, D. E. Role of alginate O acetylation in resistance of mucoid *Pseudomonas aeruginosa* to opsonic phagocytosis. Infect. Immun. 69, 1895–1901 (2001).

11. Wolfert, M. A., Roychowdhury, A. & Boons, G. J. Modification of the structure of peptidoglycan is a strategy to avoid detection by nucleotide-binding oligomerization domain protein 1. Infect. Immun. 75, 706–713 (2007).

12. Sychantha, D., Brott, A. S., Jones, C. S. & Clarke, A. J. Mechanistic pathways for peptidoglycan O-acetylation and de-O-acetylation. Front. Microbiol. 9, 1–17 (2018).

13. Bera, A., Biswas, R., Herbert, S. & Götz, F. The presence of peptidoglycan *O*-acetyltransferase in various staphylococcal species correlates with lysozyme resistance and pathogenicity. Infect. Immun. 74, 4598–4604 (2006).

14. Aubry, C., Goulard, C., Nahori, M.A., Cayet, N., Decalf, J., Sachse, M., Boneca, I.G., Cossart, P. & Dussurget, O. OatA, a peptidoglycan *O*-acetyltransferase involved in *Listeria monocytogenes* immune escape, is critical for virulence. J Infect Dis. 204, 731–40 (2011).

15. Jones, C. S., Anderson, A. C. & Clarke, A. J. Mechanism of *Staphylococcus aureus* peptidoglycan *O*-acetyltransferase A as an *O*-acyltransferase. Proc. Natl. Acad. Sci. 118, e2103602118 (2021).

16. Pearson, C. R. Tindall, S. N., Herman, R., Jenkins, H. T., Bateman, A., Thomas, G.H., Potts, J.R. & Van der Woude, M.W. Acetylation of surface carbohydrates in bacterial pathogens requires coordinated action of a two-domain membrane-bound acyltransferase. MBio 11, 1–19 (2020).

17. Weadge, J. T., Pfeffer, J.M. & Clarke, A. J. Identification of a new family of enzymes with potential *O*-acetylpeptidoglycan esterase activity in both Gram-positive and Gram-negative bacteria. BMC Microbiol. 5, 49 (2005).

18. Veyrier, F. J., Williams, A. H., Mesnage, S., Schmitt, C., Taha, M. K. & Boneca, I. G. De-O-acetylation of peptidoglycan regulates glycan chain extension and affects *in vivo* survival of *Neisseria meningitidis*. Mol. Microbiol. 87, 1100–1112 (2013).

19. Dillard, J. P. & Hackett, K. T. Mutations affecting peptidoglycan acetylation in *Neisseria gonorrhoeae* and *Neisseria meningitidis*. Infect Immun. 73, 5697–705 (2005).

20. Moynihan, P. J. & Clarke, A. J. O-Acetylated peptidoglycan: Controlling the activity of bacterial autolysins and lytic enzymes of innate immune systems. Int. J. Biochem. Cell Biol. 43, 1655–1659 (2011).

21. Moynihan, P. J. & Clarke, A. J. O-Acetylation of peptidoglycan in Gram-negative bacteria. J. Biol. Chem. 285, 13264–13273 (2010).

22. Moynihan, P. J. & Clarke, A. J. Mechanism of action of peptidoglycan *O*-acetyltransferase B involves a Ser-His-Asp catalytic triad. Biochemistry 53, 6243–6251 (2014).

23. Moynihan, P. J. & Clarke, A. J. Substrate specificity and kinetic characterization of peptidoglycan *O*-acetyltransferase B from *Neisseria gonorrhoeae*. J. Biol. Chem. 289, 16748–16760 (2014).

24. Hofmann, K. 2000. A superfamily of membrane-bound *O*-acyltransferases with implications for Wnt signaling. Trends Biochem. Sci. 25, 111–112.

25. Schultz, B. J., Snow, E. D. & Walker, S. Mechanism of D-alanine transfer to teichoic acids shows how bacteria acylate cell envelope polymers. Nat. Microbiol. 8, 1318–1329 (2023).

26. McKay Wood, B., Santa Maria, J. P., Matano, L. M., Vickery, C. R. & Walker, S. A partial reconstitution implicates DltD in catalyzing lipoteichoic acid D-alanylation. J. Biol. Chem. 293, 17985–17996 (2018).

27. Baker, P., Ricer, T., Moynihan, P. J., Kitova, E., Walvoort, M. T. C., Little, D. J., Whitney, J. C., Dawson, K., Weadge, J. T., Robinson, H., Ohman, D. E., Tipton, P., Codee, J. D. C., Klassen, J. S., Clarke, A. J. & Howell, P. L. *P. aeruginosa* SGNH hydrolase-like proteins AlgJ and AlgX have similar topology but separate and distinct roles in alginate acetylation. PLoS Pathog. 10, e1004334 (2014).

28. Franklin, M. J., Douthit, S. A. & McClure, M. A. Evidence that the algI/algJ gene cassette, required for O acetylation of *Pseudomonas aeruginosa* alginate, evolved by lateral gene transfer. J. Bacteriol. 186, 4759–4773 (2004).

29. Sychantha, D., Little, D. J., Chapman, R. N., Boons, G.-J. J., Robinson, H., Howell, P. L. & Clarke, A. J. PatB1 is an *O*-acetyltransferase that decorates secondary cell wall polysaccharides. Nat. Chem. Biol. 14, 79–85 (2018).

30. Lunderberg, J. M., Nguyen-Mau, S. M., Richter, G. S., Wang, Y. T., Dworkin, J., Missiakas, D. M. & Schneewind, O. *Bacillus anthracis* acetyltransferases PatA1 and PatA2 modify the secondary cell wall polysaccharide and affect the assembly of S-layer proteins. J. Bacteriol. 195, 977–989 (2013).

31. Spiers, A. J., Bohannon, J., Gehrig, S. M. & Rainey, P. B. Biofilm formation at the air-liquid interface by the *Pseudomonas fluorescens* SBW25 wrinkly spreader requires an acetylated form of cellulose. Mol. Microbiol. 50, 15–27 (2003).

32. Burnett, A. J. N., Rodriguez, E., Constable, S., Lowrance, B., Fish, M. & Weadge, J. T. WssI from the Gram-negative bacterial cellulose synthase is an *O*-acetyltransferase that acts on cello-oligomers with several acetyl donor substrates. J. Biol. Chem. 299, 104849 (2023).

33. Scott, W., Lowrance, B., Anderson, A. C. & Weadge, J. T. Identification of the Clostridial cellulose synthase and characterization of the cognate glycosyl hydrolase, CcsZ. PLoS One 15, e0242686 (2020).

34. Dannheim, H., Will, S. E., Schomburg, D. & Neumann-Schaal, M. *Clostridioides difficile* 630Δerm *in silico* and *in vivo* - quantitative growth and extensive polysaccharide secretion. FEBS Open Bio. 7, 602–615 (2017).

35. Brunner, J. D., Jakob, R. P., Schulze, T., Neldner, Y., Moroni, A., Thiel, G., Maier, T. & Schenck, S. Structural basis for ion selectivity in TMEM175 K+ channels. Elife 9, 1–24 (2020).

36. Ma, D., Wang, Z., Merrikh, C. N., Lang, K. S., Lu, P., Li, X., Merrikh, H., Rao, Z. & Xu. W. Crystal structure of a membrane-bound *O*-acyltransferase. Nature 562, 286–290 (2018).

37. Yang, J., Brown, M. S., Liang, G., Grishin, N. V. & Goldstein, J. L. Identification of the acyltransferase that octanoylates Ghrelin, an appetite-stimulating peptide hormone. Cell 132, 387– 396 (2008).

38. Qian, H., Zhao, X., Yan, R., Yao, X., Gao, S., Sun, X., Du, X., Yang, H., Wong, C. C. L. & Yan, N. Structural basis for catalysis and substrate specificity of human ACAT1. Nature 581, 333–338 (2020).

39. Coupland, C. E., Andrei, S. A., Ansell, T. B., Carrique, L., Kumar, P., Sefer, L., Schwab, R. A., Byrne, E. F. X., Pardon, E., Steyaert, J., Magee, A. I., Lanyon-Hogg, T., Sansom, M. S. P., Tate, E. W. & Siebold, C. Structure, mechanism, and inhibition of Hedgehog acyltransferase. Mol. Cell 81, 5025–5038.e10 (2021).

40. Pierce, M. R. & Hougland, J. L. A rising tide lifts all MBOATs: recent progress in structural and functional understanding of membrane bound *O*-acyltransferases. Front. Physiol. 14, 1–17 (2023).

41. Wang, K., Lee, C. W., Sui, X., Kim, S., Wang, S., Higgs, A. B., Baublis, A. J., Voth, G. A., Liao, M., Walther, T. C. & Farese, R. V. Jr. The structure of phosphatidylinositol remodeling MBOAT7 reveals its catalytic mechanism and enables inhibitor identification. Nat. Commun. 14, 3533 (2023).

42. Fureby, A. M., Virto, C., Adlercreutz, P. & Mattiasson, B. Acyl group migrations in 2-monoolein. Biocatal. Biotransformation 14, 89–111 (1996).

43. Serdarevich, B. Glyceride isomerizations in lipid chemistry. J. Am. Oil Chem. Soc. 44, 381–393 (1967).

44. Filip, C., Fletcher, G., Wulff, J. L. & Earhart, C. F. Solubilization of the cytoplasmic membrane of *Escherichia coli* by the ionic detergent sodium-lauryl sarcosinate. J. Bacteriol. 115, 717–722 (1973).

45. Pearson, C. R. et al. Acetylation of surface carbohydrates in bacterial pathogens requires coordinated action of a two-domain membrane-bound acyltransferase. MBio 11, e01364–20 (2020).

46. Moynihan, P. J. & Clarke, A. J. Assay for peptidoglycan *O*-acetyltransferase: A potential new antibacterial target. Anal. Biochem. 439, 73–79 (2013).

47. Anderson, A. C., Stangherlin, S., Pimentel, K. N., Weadge, J. T. & Clarke, A. J. The SGNH hydrolase family: a template for carbohydrate diversity. 32, 826–848 (2022).

48. Myers, D. K. & Kemp, A. Inhibition of esterases by the fluorides of organic acids. Nature 173, 33– 34 (1954).

49. Upton, C. & Buckley, J. T. A new family of lipolytic enzymes? Trends Biochem. Sci. 20, 178–179 (1995).

50. Mølgaard, A., Kauppinen, S. & Larsen, S. Rhamnogalacturonan acetylesterase elucidates the structure and function of a new family of hydrolases. Structure 8, 373–383 (2000).

51. Sigler, P. B., Blow, D. M., Matthews, B. W. & Henderson, R. Structure of crystalline α-chymotrypsin. J. Mol. Biol. 35, 143–164 (1968).

52. Ménard, R. & Storer, A. C. Oxyanion hole interactions in serine and cysteine proteases. Biol. Chem. Hoppe. Seyler. 373, 393–400 (1992).

53. Sychantha, D., Jones, C. S., Little, D. J., Moynihan, P. J., Robinson, H., Galley, N. F., Roper, D. I., Dowson, C. G., Howell, P. L. & Clarke, A. J. *In vitro* characterization of the antivirulence target of Gram-positive pathogens, peptidoglycan *O*-acetyltransferase A (OatA). PLoS Pathog. 13, 1–26 (2017).

54. Jones, C. S., Sychantha, D., Howell, P. L. & Clarke, A. J. Structural basis for the *O*-acetyltransferase function of the extracytoplasmic domain of OatA from *Staphylococcus aureus*. J. Biol. Chem. 295, 8204–8213 (2020).

55. Evans, R., O’Neill, M., Pritzel, A., Antropova, N., Senior, A., Green, T., Žídek, A., Bates, R., Blackwell, S., Yim, J., Ronneberger, O., Bodenstein, S., Zielinski, M., Bridgland, A., Potapenko, A., Cowie, A., Tunyasuvunakool, K., Jain, R., Clancy, E., Kohli, P., Jumper, J., Hassabis. D. Protein complex prediction with AlphaFold-Multimer. bioRxiv 10.1101/2021.10.04.463034 (2022).

56. Sui, X., Wang, K., Gluchowski, N. L., Elliott, S.D., Liao, M., Walther, T.C. & Farese, R. V. Structure and catalytic mechanism of a human triacylglycerol-synthesis enzyme. Nature 581, 323–328 (2020).

57. Jiang, Y., Benz, T. L. & Long, S.B. Substrate and product complexes reveal mechanisms of Hedgehog acylation by HHAT. Science 372, 1215–1219 (2021).

58. Hunter, T. Tyrosine phosphorylation: thirty years and counting. Curr. Opin. Cell Biol. 21,140–146 (2009).

59. Grangeasse, C., Cozzone, A. J., Deutscher, J. & Mijakovic, I. Tyrosine phosphorylation: an emerging regulatory device of bacterial physiology. Trends Biochem. Sci. 32, 86–94 (2007).

60. Moore, K. L. The biology and enzymology of protein tyrosine O-sulfation. J. Biol. Chem. 278, 24243–24246 (2003).

61. Tse, Y. C., Kirkegaard, K. & Wang, J. C. Covalent bonds between protein and DNA. Formation of phosphotyrosine linkage between certain DNA topoisomerases and DNA. J. Biol. Chem. 255, 5560–5565 (1980).

62. Redinbo, M. R., Stewart, L., Kuhn, P., Champoux, J. J. & Hol, W. G. J. Crystal structures of human topoisomerase I in covalent and noncovalent complexes with DNA. Science 279, 1504–1513 (1998).

63. Cheng, C., Kussie, P., Pavletich, N. & Shuman, S. Conservation of structure and mechanism between eukaryotic topoisomerase I and site-specific recombinases. Cell 92, 841–850 (1998).

64. Meinke, G., Bohm, A., Hauber, J., Pisabarro, M. T. & Buchholz, F. Cre recombinase and other tyrosine recombinases. Chem. Rev. 116,12785–12820 (2016).

65. Watts, A. G., Damager, I., Amaya, M. L., Buschiazzo, A., Alzari, P., Frasch, A. C. & Withers, S. G. *Trypanosoma cruzi* trans-sialidase operates through a covalent sialyl-enzyme intermediate: Tyrosine is the catalytic nucleophile. J. Am. Chem. Soc 125, 7532–7533 (2003).

66. Vavricka, C. J., Liu, Y., Kiyota, H., Sriwilaijaroen, N., Qi, J., Tanaka, K., Wu, Y., Li, Q., Li, Y., Yan, J., Suzuki, Y. & Gao, G. F. Influenza neuraminidase operates via a nucleophilic mechanism and can be targeted by covalent inhibitors. Nat. Commun. 4,1491 (2013).

67. Kim, J. H., Resende, R., Wennekes, T., Chen, H. M., Bance, N., Buchini, S., Watts, A. G., Pilling, P., Streltsov, V. A., Petric, M., Liggins, R., Barrett, S., McKimm-Breschkin, J. L., Niikura, M. & Withers, S. G. (2013) Mechanism-based covalent neuraminidase inhibitors with broad-spectrum influenza antiviral activity. Science 340, 71–75.

68. O’Neil, M. J., Heckelman, P. E., Dobbelaar, P. H. & Roman, K. J. eds. The Merck Index (Royal Society of Chemistry) 15th Ed. (2013).

69. Dawson, R. M. C., Elliott, D. C., Elliott, W. H. & Jones, K. M. Data for Biochemical Research (Clarendon Press) 3rd Ed (1989).

70. Clarke, A. J. & Dupont, C. O-Acetylated peptidoglycan: Its occurrence, pathobiological significance, and biosynthesis. Can. J. Microbiol. 38, 85–91 (1992).

71. Lear, A. L., & Perkins, H, J. O-Acetylation of peptidoglycan in *Neisseria gonorrhoeae*. Investigation of lipid-linked intermediates and glycan chains newly incorporated into the cell wall. J. Gen. Microbiol. 132, 2413–1420 (1986)

72. Williams, A. H., Veyrier, F. J., Bonis, M., Michaud, Y., Raynal, B., Taha, M. K., White, S. W., Haouz, A., and Boneca, I. G. Visualization of a substrate-induced productive conformation of the catalytic triad of the *Neisseria meningitidis* peptidoglycan *O*-acetylesterase reveals mechanistic conservation in SGNH esterase family members. Acta Crystallogr. Sect. D Biol. Crystallogr. 70, 2631–2639 (2014).

73. Liu, H. & Naismith, J. H. An efficient one-step site-directed deletion, insertion, single and multiple-site plasmid mutagenesis protocol. BMC Biotechnol. 8, (2008).

74. Inoue, H., Nojima, H. & Okayama, H. High efficiency transformation of *Escherichia coli* with plasmids. Gene 96, 23–28 (1990).

75. Brott, A. S., Jones, C. S. & Clarke, A. J. Development of a high throughput screen for the identification of inhibitors of peptidoglycan *O*-acetyltransferases, new potential antibacterial targets. Antibiotics 8, 65 (2019).

76. Reverter, D. & Lima, C. D. Preparation of SUMO proteases and kinetic analysis using endogenous substrates. in SUMO Protocols (ed. Ulrich, H. D.) pp. 225–240 (Humana Press, New York, 2009).

77. Neu, H. C. & Heppel, L. A. The release of enzymes from *Escherichia coli* by osmotic shock and during the formation of spheroplasts. J. Biol. Chem. 240, 3685–3692 (1965).

78. Kabsch, W. XDS. Acta Crystallogr. Sect. D Biol. Crystallogr. 66, 125–132 (2010).

79. Adams, P. D. et al. PHENIX: A comprehensive Python-based system for macromolecular structure solution. Acta Crystallogr. Sect. D Biol. Crystallogr. 66, 213–221 (2010).

80. Emsley, P., Lohkamp, B., Scott, W. G. & Cowtan, K. Features and development of Coot. Acta Crystallogr. Sect. D Biol. Crystallogr. 66, 486–501 (2010).

81. Teufel, F., Almagro Armenteros, J. J., Johansen, A. R., Gíslason, M. H., Pihl, S. I., Tsirigos, K. D., Winther, O., Brunak, S., von Heijne, G. & Nielsen, H. SignalP 6.0 predicts all five types of signal peptides using protein language models. Nat. Biotechnol. 40, 1023–1025 (2022).

82. Krogh, A., Larsson, B., Von Heijne, G. & Sonnhammer, E. L. L. Predicting transmembrane protein topology with a hidden Markov model: Application to complete genomes. J. Mol. Biol. 305, 567– 580 (2001).

