## Supporting Information for "The mechanism of peptidoglycan O-acetylation in Gram-negative bacteria typifies bacterial MBOAT-SGNH acyltransferases"

[†]These authors contributed equally

**This file includes:**

- Tables S1 to S2
- Figures S1 to S15
- SI References

### Table S1. X-ray data collection and refinement statistics.

| Dataset | ***Ng*PatB$_{\Delta100}$ SeMet** | ***Ng*PatB$_{\Delta100}$ Native** | ***Ng*PatB$_{\Delta100}$ MeS** | ***Cj*PatB$_{\Delta113}$ Native** |
|---|---|---|---|---|
| **Data Collection** | | | | |
| Beamline | CLS 08ID-1 | CLS 08ID-1 | CLS 08ID-1 | CLS 08ID-1 |
| Wavelength (Å) | 0.97828 | 0.97949 | 0.97828 | 0.97949 |
| Space group | $P\ 3_2$ | $P\ 3_2$ | $P\ 3_2$ | $P\ 6_1$ |
| Unit cell parameters (Å/°) | $a = b = 144.5$ | $a = b = 144.9$ | $a = b = 145.7$ | $a = b = 55.83$ |
| | $c = 77.95$ | $c = 80.14$ | $c = 78.41$ | $c = 136.44$ |
| | $\alpha = \beta = 90$ | $\alpha = \beta = 90$ | $\alpha = \beta = 90$ | $\alpha = \beta = 90$ |
| | $\gamma = 120$ | $\gamma = 120$ | $\gamma = 120$ | $\gamma = 120$ |
| Resolution range (last shell) | 49.21 – 1.2 | 41.83 – 1.19 | 49.15 – 1.43 | 48.35 – 1.90 |
| (Å) | (1.24 – 1.20) | (1.22 – 1.19) | (1.47 – 1.43) | (1.968 – 1.90) |
| | 485015 | 1075309 | 534809 | 386275 |
| Total number of reflections | 79513 | 102048 | 56726 | 18944 |
| Number of unique reflections | 6.1 | 10.5 | 32 | 20.4 |
| Redundancy | 99.03 (11.80) | 99.96 (94.6) | 99.99 (43.0) | 99.98 (100.00) |
| Completeness (last shell) (%) | 4.32 (0.27) | 17.4 (1.3) | 2.8 (1.24) | 28.39 (9.23) |
| Average $I/\sigma$ ($I$) (last shell) | 9.7 (405) | 7.7 (301) | 6.2 (186) | 9.9 (71) |
| $R_{merge}$ (last shell) (%)[1] | 0.998 (0.091) | 0.999 (0.203) | 0.996 (0.440) | 0.999 (0.977) |
| $CC_{1/2}$ (last shell)[2] | | | | |
| **Refinement** | | | | |
| Resolution range (Å) | 49.21 – 1.50 | 41.83 – 1.30 | 49.15 – 1.8 | 39.45 – 1.9 |
| $R_{work}/R_{free}$ (%)[2] | 20.4 / 22.2 | 16.6 / 18.9 | 19.6 / 22.9 | 16.8 / 20.4 |
| Number of atoms | 1909 | 1991 | 1890 | 1857 |
| Protein | 1758 | 1782 | 1754 | 1753 |
| Water | 151 | 204 | 132 | 103 |
| Ligand | 0 | 5 | 4 | 1 |
| Average B-factor (Å$^2$)[3] | 18.9 | 24.5 | 26.2 | 21.7 |
| Protein | 18.3 | 23.6 | 25.6 | 24.1 |
| Water | 25.7 | 32.3 | 32.7 | 30.4 |
| Ligand | N/A | 47.8 | 37.7 | 30.0 |
| RMS Bond lengths (Å) | 0.006 | 0.005 | 0.006 | 0.007 |
| RMS Bond angles (°) | 0.805 | 0.764 | 0.711 | 0.770 |
| Ramachandran favoured (%) | 97.21 | 97.27 | 97.21 | 97.17 |
| Ramachandran allowed (%) | 2.79 | 2.73 | 2.79 | 2.83 |
| PDB accession ID | 7TLV | 7TJB | 7TRR | 8TLB |

Values in parentheses correspond to the highest resolution shell.

*I/σ(I):* intensity of a group of reflections divided by the standard deviation of those reflections.

[1]$R_{merge} = \sum\sum |I(k) - \langle I \rangle| / \sum I(k)$, where $I$(k) and $\langle I \rangle$ represent the diffraction intensity values of the individual measurements and the corresponding mean values. The summation is over all unique measurements.

[2]$R_{work} = \sum ||F_{obs}| - k|F_{calc}|| / |F_{obs}|$, where $F_{obs}$ and $F_{calc}$ are the observed and calculated structure factors, respectively. $R_{free}$ is the sum extended over a subset of reflections excluded from all stages of the refinement.

[3]As calculated using MolProbity (1)

**Table S2: Strains, plasmids, and primers used in this study.**

| **Strains** | Description or characteristics | Source or reference |
|---|---|---|
| *C. jejuni* 81-176 | Wild type *C. jejuni* strain isolated from a diarrheic patient | 2 |
| *E. coli* DH5α | Laboratory strain used for cloning: F– φ80*lac* ZΔM15 Δ(*lac* ZYA-*arg* F)U169 *rec* A1 *end*A1 *hsd*R17(rK–, mK+) *pho* A *sup* E44 λ–thi-1 *gyr*A96 *rel* A1 | Thermo Fisher |
| *E. coli* C43 (DE3) | Laboratory strain used for overexpression: F–*omp* T *hsd*SB (rB–, mB–) *gal dcm* (DE3), contains further mutations for overexpression of toxic proteins | 3 |
| *E. coli* BL21 (DE3) | Laboratory strain used for overexpression: F–*omp* T *hsd*SB (rB–, mB–) *gal dcm rne* 131 (DE3) | Invitrogen |
| *E. coli* Stellar | *F- endA1 supE44 thi-1 recA1 relA1 gyrA96 phoA Φ80dlacZΔM15 Δ(lacZYA-argF) U169 Δ*(*mrr-hsdRMS-mcrBC*) *ΔmcrA λ-* | Takara Bio |
| *N. gonorrhoeae* FA1090 | Serum-resistant, proline-requiring strain isolated from a patient with probable disseminated gonococcal infection | 4 |
| **Plasmids** | | |
| pBAD | T7-promoter driven *E. coli* vector with *araBAD* promoter for N-terminal $His_6$-fusion proteins | Invitrogen |
| pET28a(+) | T7-promoter driven *E. coli* expression vector for $His_6$-fusion proteins | Novagen |
| pET26b(+) | T7-promoter driven *E. coli* expression vector with *pelB* signal sequence for periplasmic localization of $His_6$-fusion proteins | Novagen |
| pACAB7 | pBAD HisA expression plasmid containing $His_6$-SUMO tagged *Ng*$PatB_{\Delta100}$ | 5 |
| pACAA1 | pET28a expression plasmid containing C-terminally $His_6$ tagged *Cj*$PatB_{\Delta31}$ | This study |
| pACAA2 | pET28a expression plasmid containing C-terminally $His_6$ tagged *Cj*$PatB_{\Delta113}$ | This study |
| pACAA3 | pET28a expression plasmid containing C-terminally $His_6$ tagged *Cj*PatB | This study |
| pACAA4 | pET28a expression plasmid containing N-terminally $His_6$ tagged *Cj*$PatB_{\Delta31}$ | This study |
| pACAA5 | pET28a expression plasmid containing N-terminally $His_6$ tagged *Cj*$PatB_{\Delta113}$ | This study |
| pBS216[1] | pET-26b(+) derivative for expression of *Cj*PatA tandem fusion protein | This study |
| pJL002 | pBS216 derivative for expression of *Cj*$PatA^{Y455F}$ | This study |
| pJL005 | pBS216 derivative for expression of *Cj*$PatA^{H315A}$ | This study |

**Table S1: Strains, plasmids, and primers used in this study - *continued***

| **Primers** | | |
| --- | --- | --- |
| *Cj*PatB$_{\Delta 31}$ NdeI Fwd | CTAGCATATGCAAAAATATCACTTTGCATTCTATCC | IDT[2], this study |
| *Cj*PatB XhoI Rev | ATCGCTCGAGTTTACTAGCATTTTCTTCCTTAAGTTTT ATGTG | IDT, this study |
| *Cj*PatB FL CHis NcoI Fwd | ACGTCCATGGGCAGTGTAGTTAGATTTTTCTTTATTTT GATTATAGTG | IDT, this study |
| *Cj*PatB(Δ113) NdeI Fwd | CTAGCATATGGATGCAAATATTAGTTTTATCGAC | IDT, this study |
| *Cj*PatB XhoI + stop Rev (N-His) | ATCGCTCGAGTTATTTACTAGCATTTTCTTCCTTAAGT TTTATGTG | IDT, this study |
| ABP16 Fwd | /5Phos/GGCACCGAATGGAAACAGGGC | IDT, this study |
| APB6r Rev | /5Phos/ACCACCAATCTGTTCTCTGTGAGCC | IDT, this study |
| oBS584 | AATTTTGTTTAACTTTAAGAAGGAGATATACATATGAA ATACCTG | This study |
| oBS585 | AGCCGGATCTCAGTGGTGG | This study |
| oJL003 | CTCAAGCTTTTAATAACTCGAGCAC | This study |
| oJL004 | AAAATAAAATCCGGAATGCCATCTG | This study |
| oJL008 | GGGGAACACGCTGGCGTTTATTG | This study |
| oJL009 | GCCCACATGCCTGACAGGATAAACG | This study |

[1]pBS216 encodes ss$_{PelB}$-linker-His$_{10}$-linker-MBP-linker-[HRV 3C protease cleavage site]-linker **Cj*PatA$^{M1I}$, where ss$_{PelB}$ = signal sequence of *Erwinia carotovora* pectate lyase B, MBP = *E. coli* maltose-binding protein *Note: the *Campylobacter jejuni* PatA sequence was "codon-harmonized" for expression in *E. coli* using the Codon Harmonizer Galaxy server, (7)
[2]IDT, Integrated DNA Technologies

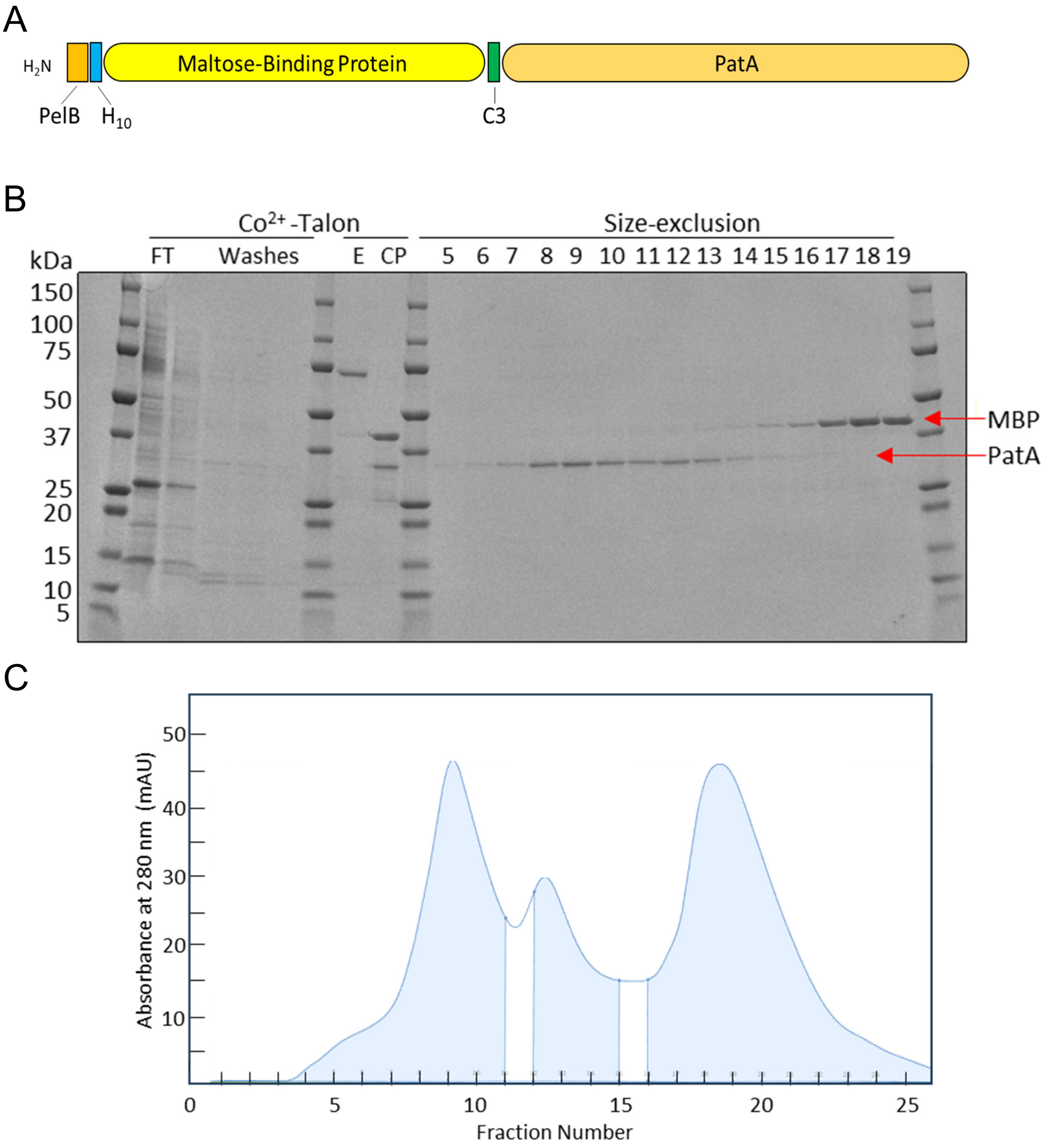


**Fig S1**. **Construct design and purification of recombinant PatA.** (A) The PatA expression construct design involving an N-terminal leader sequence (22 residues) from the pectate lyase gene *pelB* from *Erwinia caratovora*, followed by a $His_{10}$ tag, the *E. coli* maltose binding protein (MBP) lacking its N-terminal signal sequence, and a 3C protease recognition site (LEVLFQ/GP) fused in tandem to the 5' end of *patA.* (B) Representative SDS-PAGE analysis of PatA purification by $Co^{2+}$-Talon affinity and size-exclusion chromatographies. FT, column flow-through; E, column elution; CP, cleavage products of 3C protease. Lanes at right labelled 5 through 19 correspond to the respective numbered fractions of the chromatogram below. (C) Representative chromatogram of fraction labelled CP in panel B subjected to size-exclusion chromatography.

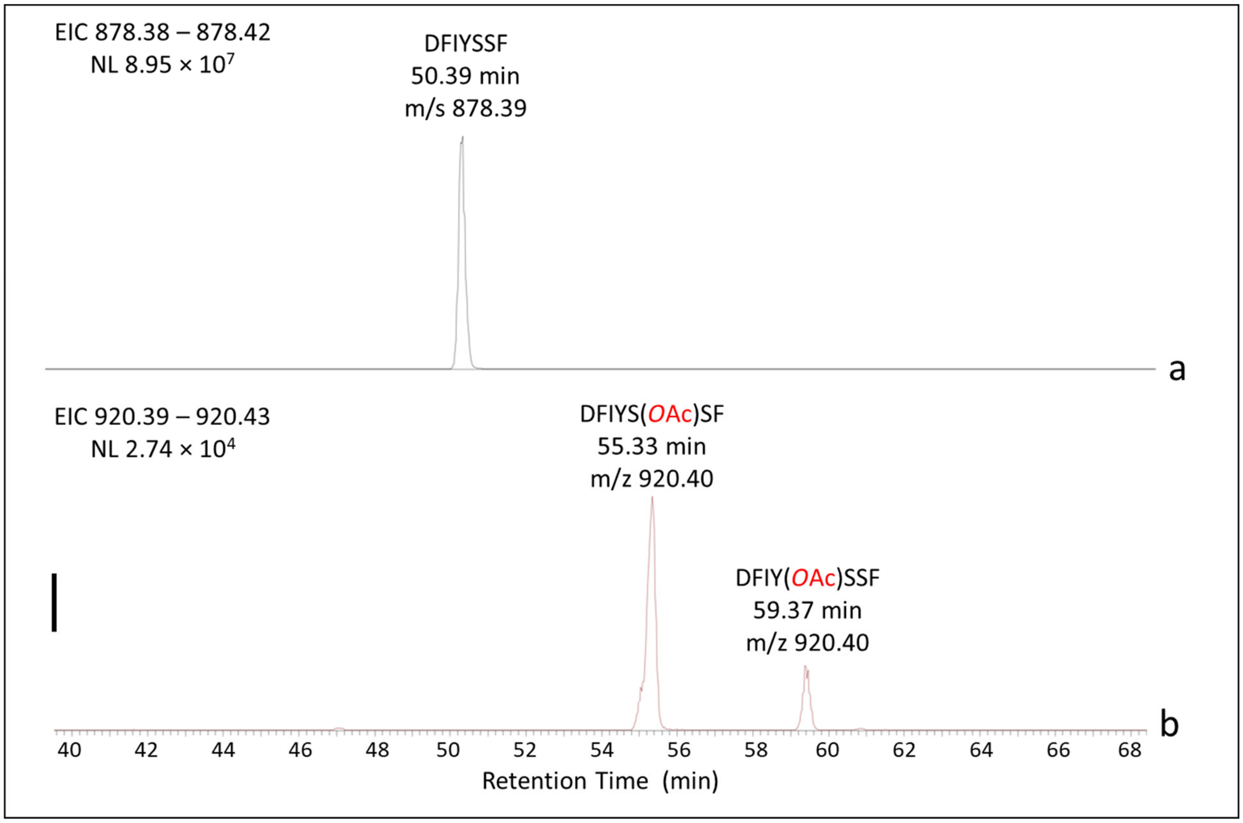


**Fig. S2**. **Trapping of acetylated-PatA intermediate**. LC-MS analysis of AspN-digested PatA following its incubation with acetyl-CoA. Extracted ion chromatograms (EIC) of the (a) native and (b) acetylated DFIYSSF peptides. The solid bar on the left denotes 25% absorbance at 210 nm relative to largest peak.

**Predicted Fragmentation Pattern**

| Seq | # | b: Δ Error | b | y | y: Δ Error | +1 |
|---|---|---|---|---|---|---|
| D | 1 | --- | 116.034 | --- | --- | 7 |
| F | 2 | -0.241 | 263.103 | 763.366 | --- | 6 |
| I | 3 | -0.137 | 376.187 | 616.298 | -0.356 | 5 |
| Y | 4 | -1.066 | 539.250 | 503.214 | 0.329 | 4 |
| S | 5 | -6.546 | 626.282 | 340.150 | 0.051 | 3 |
| S | 6 | --- | 713.314 | 253.118 | 0.069 | 2 |
| F | 7 | --- | --- | 166.086 | 0.291 | 1 |

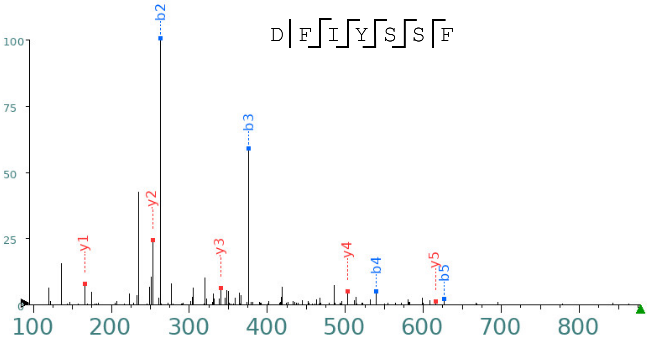


**Predicted Fragmentation Pattern**

| Seq | # | b: Δ Error | b | y | y: Δ Error | +1 |
|---|---|---|---|---|---|---|
| D | 1 | --- | 116.034 | --- | --- | 7 |
| F | 2 | 0.107 | 263.103 | 805.377 | --- | 6 |
| I | 3 | 0.107 | 376.187 | 658.308 | 2.347 | 5 |
| Y# | 4 | -1.314 | 581.261 | 545.224 | 0.853 | 4 |
| S | 5 | 3.995 | 668.293 | 340.150 | -3.538 | 3 |
| S | 6 | --- | 755.325 | 253.118 | 0.672 | 2 |
| F | 7 | --- | --- | 166.086 | 0.658 | 1 |

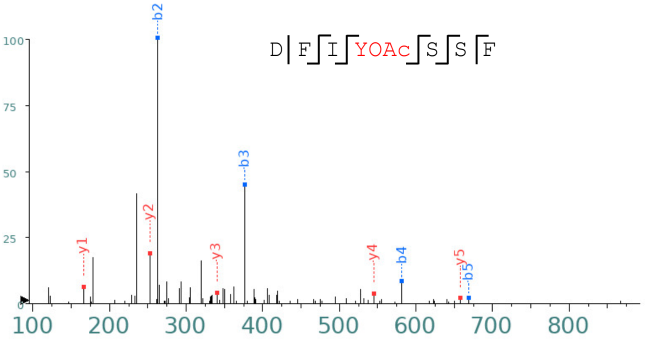


**Predicted Fragmentation Pattern**

| Seq | # | b: Δ Error | b | y | y: Δ Error | +1 |
|---|---|---|---|---|---|---|
| D | 1 | --- | 116.034 | --- | --- | 7 |
| F | 2 | 0.339 | 263.103 | 805.377 | --- | 6 |
| I | 3 | -0.056 | 376.187 | 658.308 | --- | 5 |
| Y | 4 | -0.953 | 539.250 | 545.224 | 1.189 | 4 |
| S# | 5 | --- | 668.293 | 382.161 | --- | 3 |
| S | 6 | --- | 755.325 | 253.118 | -0.172 | 2 |
| F | 7 | --- | --- | 166.086 | 1.577 | 1 |

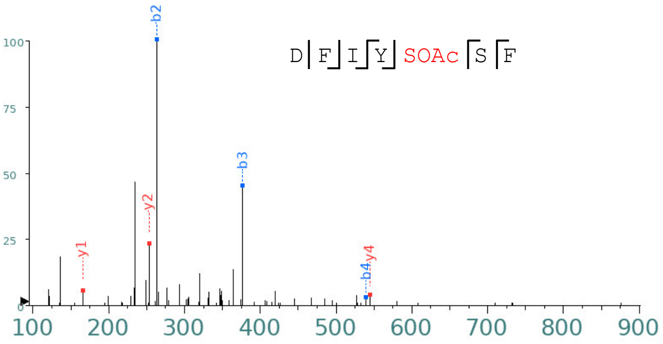


**Fig. S3**. **Representative MS/MS analysis of the *O*-acetylated PatA peptides**. Fragment ion table (left) and MS/MS spectra (right) for the peptides as identified.

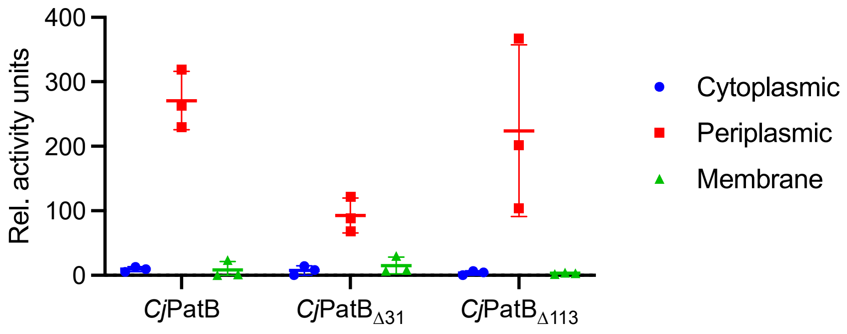


**Fig. S4. Cellular fractionation controls.** Relative alkaline phosphatase activity for each of the isolated subcellular fractions expressing *Cj*PatB forms depicted in Figure 2.

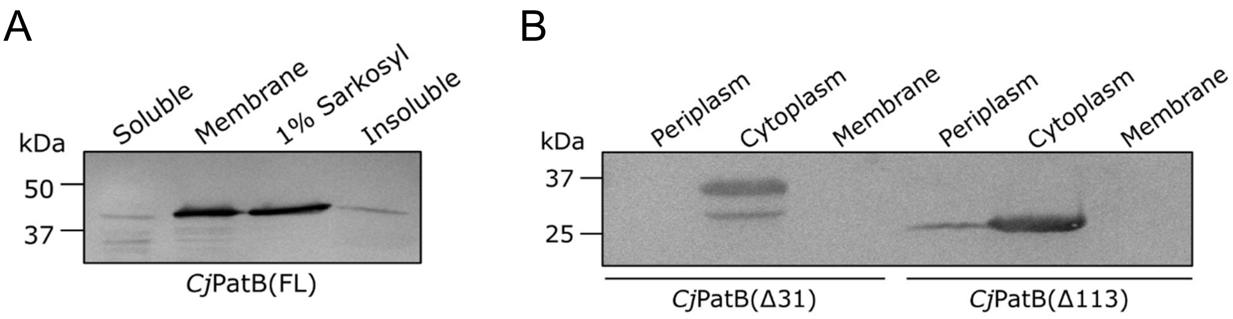


**Fig. S5. Localization of *Cj*PatB and engineered variants**. (A) Anti-His$_6$ Western blot analysis of subcellular fractions of full-length recombinant *Cj*PatB isolated from 18 h cultures of *E. coli* BL21 transformed with pACAA3. Sarkosyl (sodium lauryl sarkosinate) is a detergent selective for solubilization of the cytoplasmic membrane. (B) Localization of truncated recombinant forms of *Cj*PatB lacking either the predicted N-terminal TM helix (Δ31) or the entire N-terminal domain (Δ113).

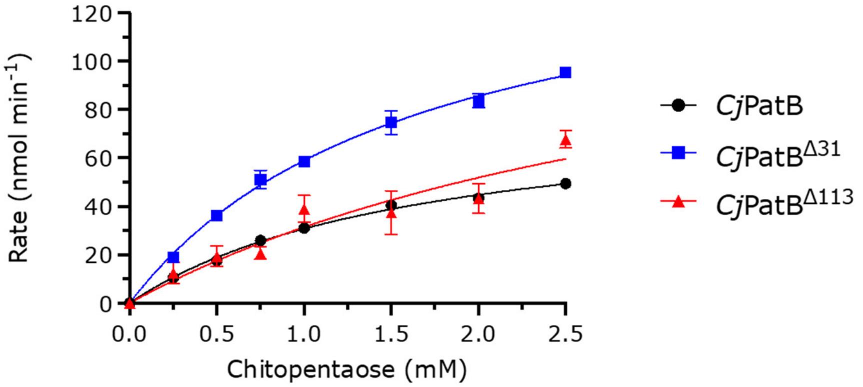


**Fig. S6. Activity of PatB as an *O*-acetyltransferase.** Dependence of the enzyme variants (1 μM) in 50 mM sodium phosphate pH 7.0 on chitopentaose concentrations as acceptor with 2 mM *p*NP-Ac as acetyl donor.

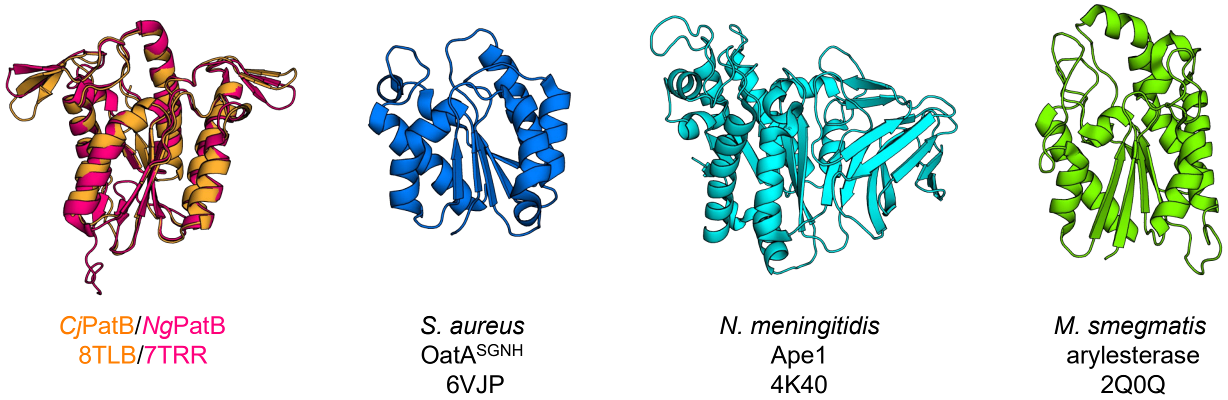


**Fig. S7. Comparison of the SGNH hydrolase domain of PatB with those of other family members.** The structures of PatB from *C. jejuni* (orange) and *N. gonorrhoeae* (magenta) contain the central α/β hydrolase fold characteristic of the SGNH hydrolases but in addition possess unique β-hairpin motifs not seen in other family members

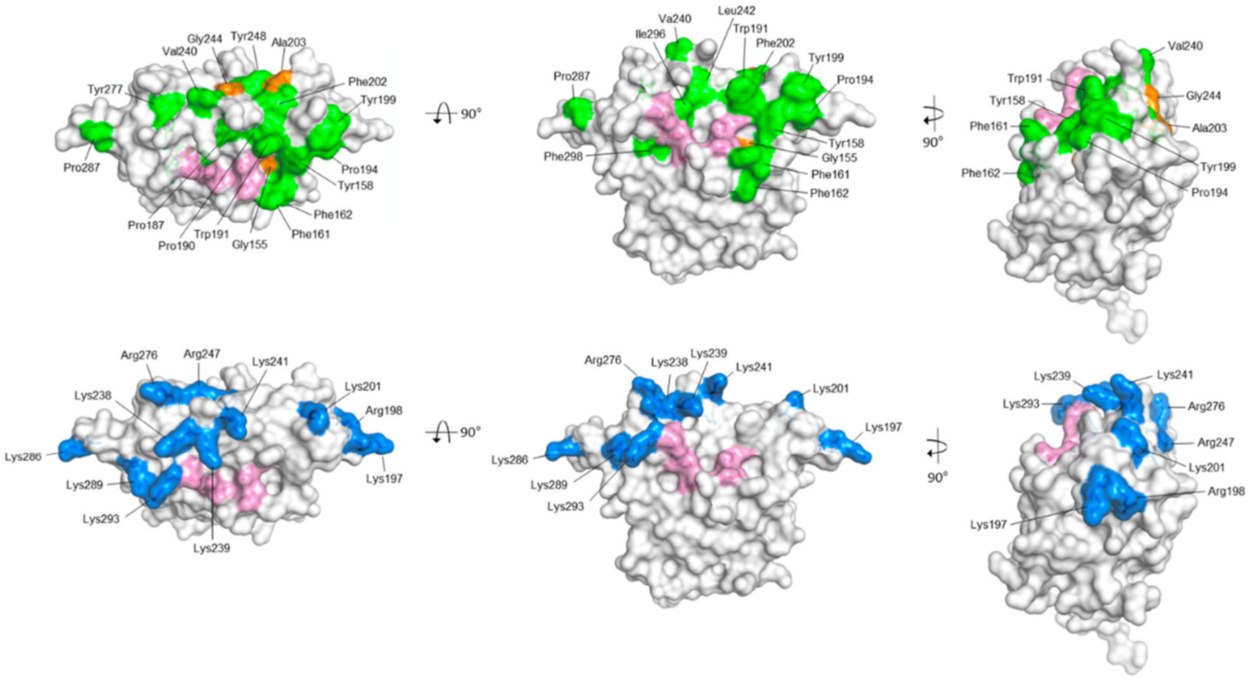


**Fig. S8. Surface non-polar and charged residues in PatB.** Surface-exposed non-polar (top row) and positively-charge (bottom row) residues in *Ng*PatB$_{\Delta 100}$ are depicted in green and blue, respectively; the catalytic and oxyanion hole residues are in pink.

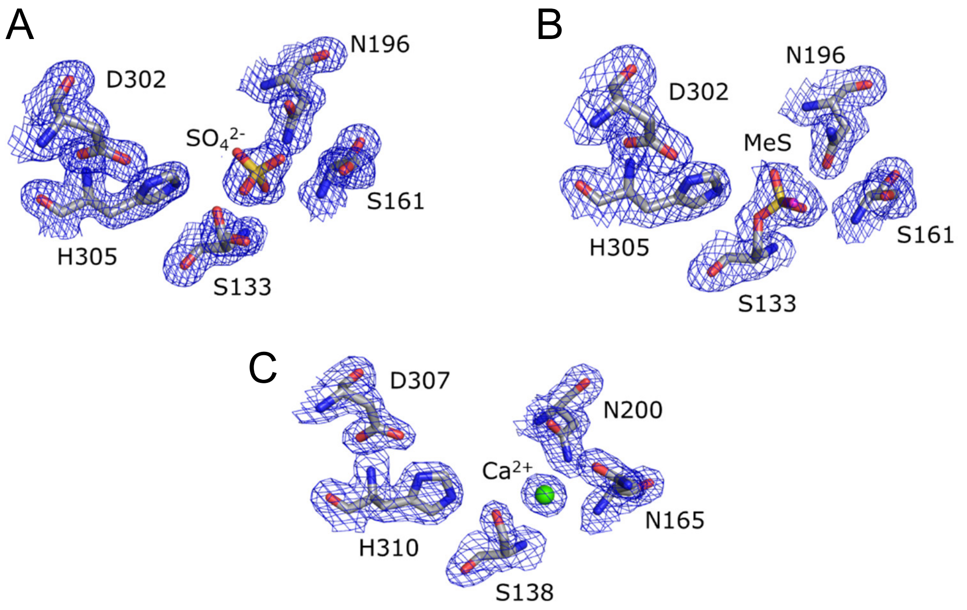


**Fig. S9. Accessory electron density in the PatB active sites.** (A) The native structure of *Ng*PatB (7TJB) contains a tetrahedral electron density adjacent to Ser133 that can be attributed to a sulfate anion. Two discrete conformers of the Ser133 sidechain are observed oriented both toward and away from the His305 sidechain. (B) The MS-bound structure of *Ng*PatB (7TRR) contains a similar tetrahedral electron density, but this density is continuous with the Ser133 sidechain and unambiguously indicates a covalent MS adduct. (C) The native structure of *Cj*PatB (8TLB) displays only one conformer of the catalytic Ser138. A $Ca^{2+}$ ion is observed occupying the same position as sulfate in the *Ng*PatB structure. All maps shown represent the 2Fo-Fc map contoured at 2σ.

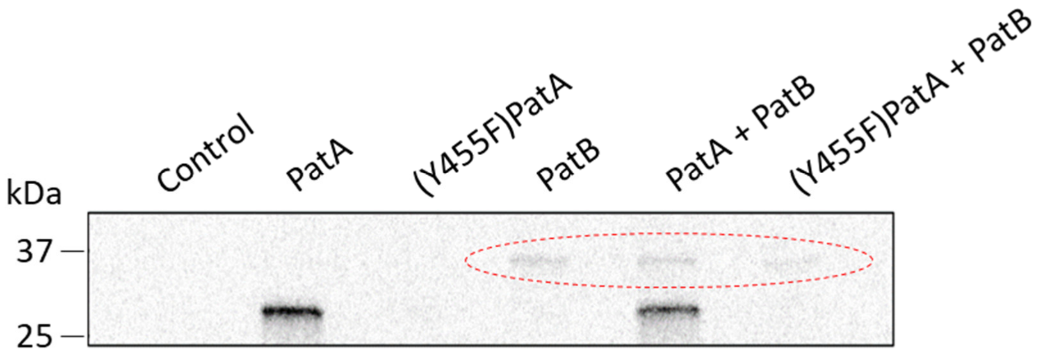


**Fig. S10. Detection of acetyl-transfer from PatA to PatB.** The enzymes (5 µM) in 50 mM HEPES, pH 7.4 containing 150 mM NaCl were incubated with 100 µM [1,2-$^{14}$C]acetyl-CoA for 15 min prior to SDS-PAGE with autoradiography detection. The dashed red circle denotes the weak signal detected from PatB in both the absence and presence of PatA.

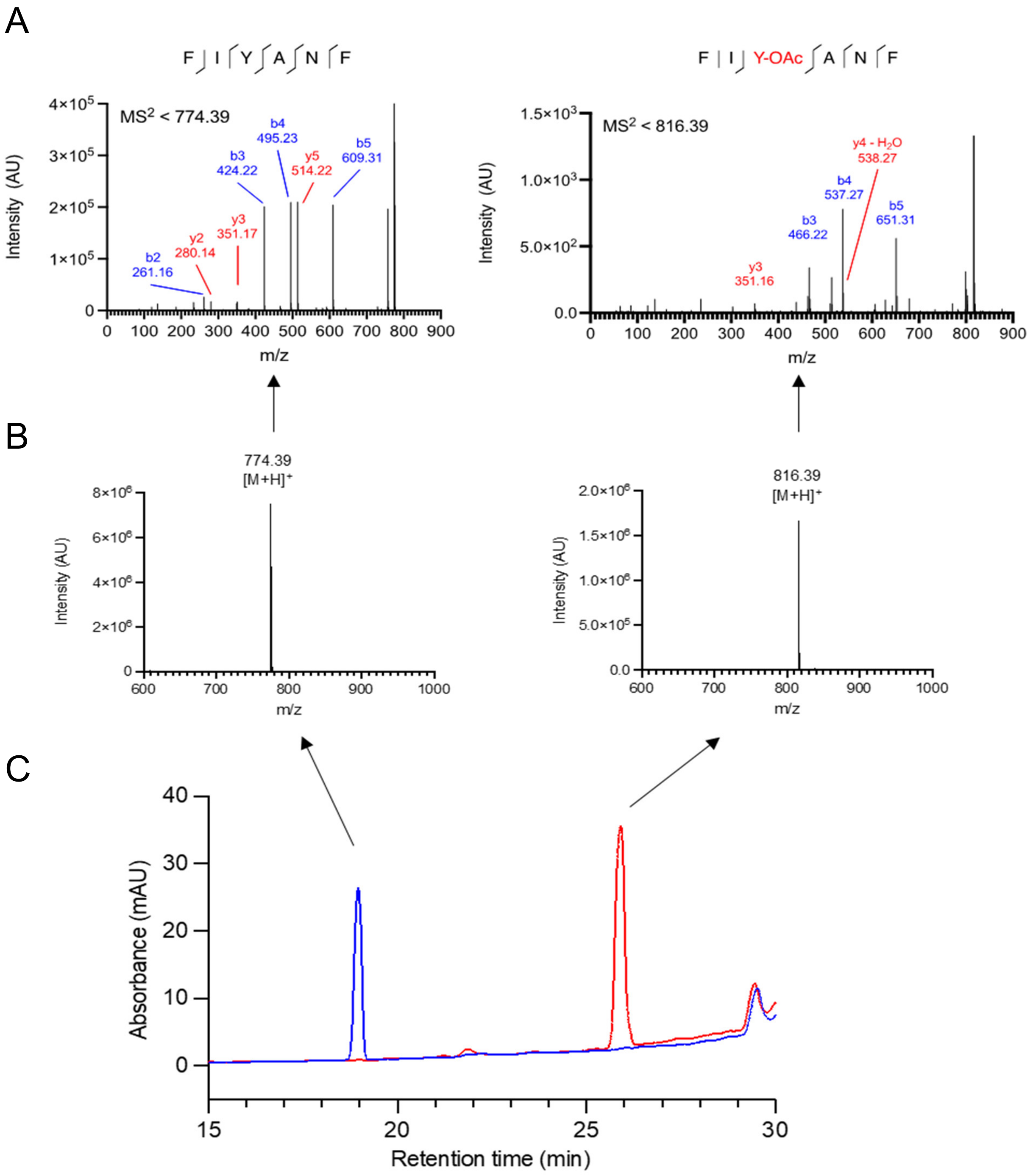


**Fig. S11**. **Purification and assignment of the synthetic peptide product**. (A) Representative HPLC chromatogram of the starting material (1) and acetylated product (2) demonstrates stoichiometric acetylation of the peptide. (B) MS identification matches the expected size of the starting material (1), 773.4 Da and confirms the product is singly acetylated (2), 815.4 Da. (C) MS/MS fragmentation allows the assignment of the 42.0 Da acetyl group to the Tyr residue of the peptide.

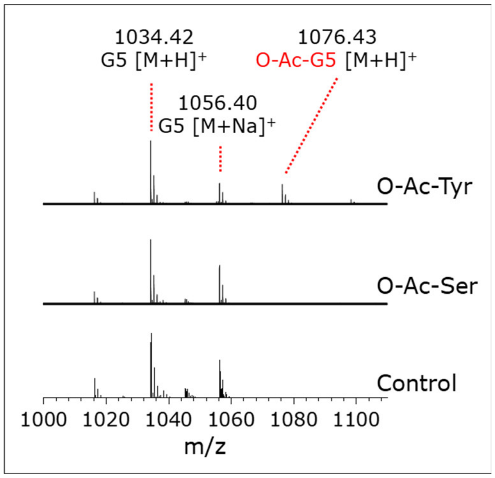


**Fig. S12**. ***O*-Acety-tyrosine as a substrate for PatB.** Representative LC-MS analysis of reaction products of PatB acting as an *O*-acetyltransferase. *Cj*PatB (1 µM) in 50 mM sodium phosphate pH 7.0 was incubated with 5 mM chitopentaose as acceptor in the absence (control) or presence of 1 mM *O*-acetyl-Ser or *O*-acetyl-Tyr.

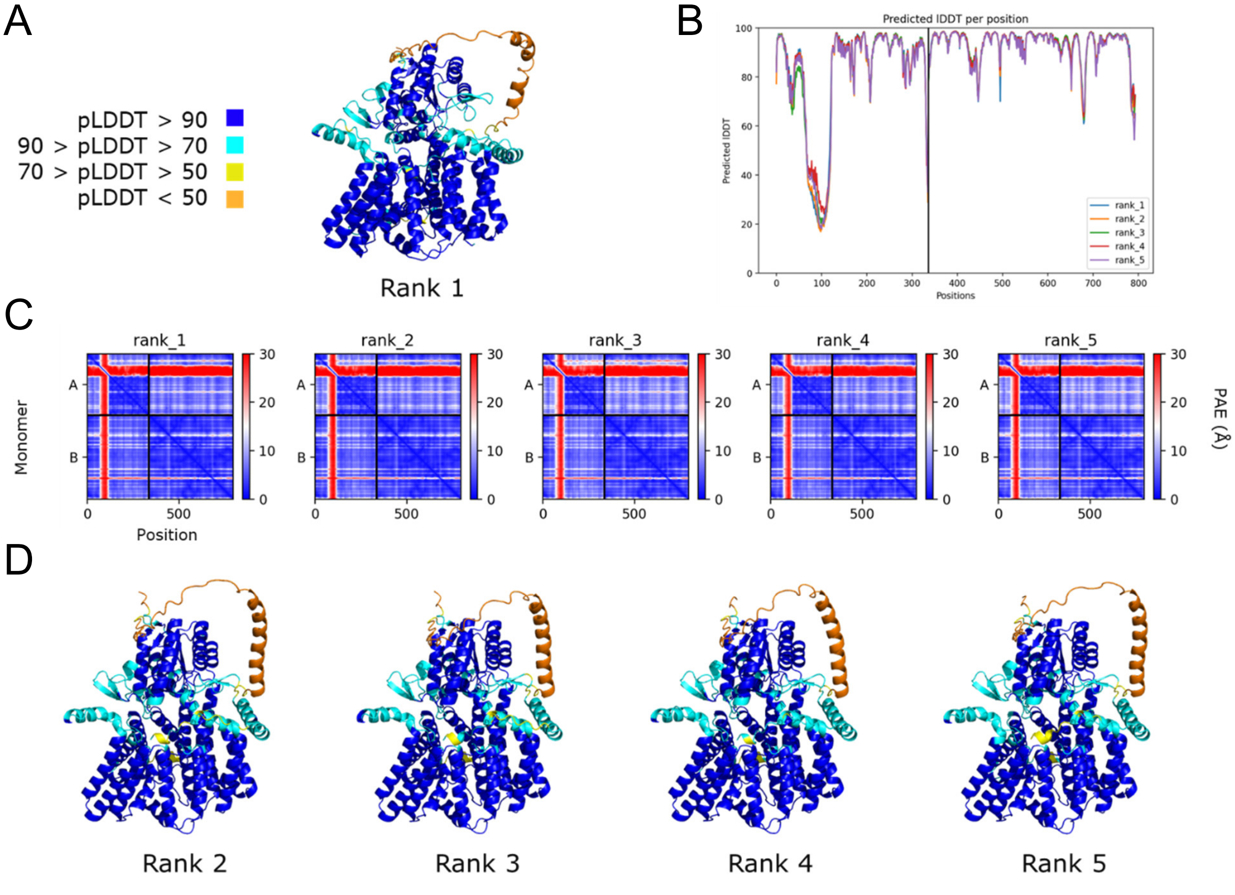


**Fig. S13**. **AlphaFold Multimer model of *C. jejuni* PatAB**. (A) The top-scoring model of PatAB. The model is coloured by predicted local distance difference test (pLDDT) values. Model confidence is based upon these scores which are quantized into very high (pLDDT > 90; blue), confident (90 > pLDDT > 70; cyan), low (70 > pLDDT > 50; yellow), and very low (50 > pLDDT; orange). (B) pLDDT scores as a function of residue for each of the top five scoring models. (C) The predicted-aligned error (PAE) heatmap for each of the top scoring models. Each x,y value represents the predicted coordinate error of residue x (in Å) between the true and predicted structures if they were aligned on position y. Monomer A is PatB and monomer B is PatA. (D) The 2$^{nd}$ to 5$^{th}$ ranked AlphaFold Multimer models of PatAB coloured by pLDDT as in panel A.

ATGAAATACCTGCTGCCGACCGCTGCTGCTGGTCTGCTGCTCCTCGCTGCCCAGCCGGCGATGG
CCGCAGATAGCGGTAGCTCTGGCCACCATCATCACCATCACCATCACCATCATGGCAGCTCTGG
TAGCAAAATCGAAGAAGGTAAACTGGTAATCTGGATTAACGGCGATAAAGGCTATAACGGTCTC
GCTGAAGTCGGTAAGAAATTCGAGAAAGATACCGGAATTAAAGTCACCGTTGAGCATCCGGATA
AACTGGAAGAGAAATTCCCACAGGTTGCGGCAACTGGCGATGGCCCTGACATTATCTTCTGGGC
ACACGACCGCTTTGGTGGCTACGCTCAATCTGGCCTGTTGGCTGAAATCACCCCGGACAAAGCG
TTCCAGGACAAGCTGTATCCGTTTACCTGGGATGCCGTACGTTACAACGGCAAGCTGATTGCTT
ACCCGATCGCTGTTGAAGCGTTATCGCTGATTTATAACAAAGATCTGCTGCCGAACCCGCCAAA
AACCTGGGAAGAGATCCCGGCGCTGGATAAAGAACTGAAAGCGAAAGGTAAGAGCGCGCTGATG
TTCAACCTGCAAGAACCGTACTTCACCTGGCCGCTGATTGCTGCTGACGGGGGTTATGCGTTCA
AGTATGAAAACGGCAAGTACGACATTAAAGACGTGGGCGTGGATAACGCTGGCGCGAAAGCGGG
TCTGACCTTCCTGGTTGACCTGATTAAAAACAAACACATGAATGCAGACACCGATTACTCCATC
GCAGAAGCTGCCTTTAATAAAGGCGAAACAGCGATGACCATCAACGGCCCGTGGGCATGGTCCA
ACATCGACACCAGCAAAGTGAATTATGGTGTAACGGTACTGCCGACCTTCAAGGGTCAACCATC
CAAACCGTTCGTTGGCGTGCTGAGCGCAGGTATTAACGCCGCCAGTCCGAACAAAGAGCTGGCA
AAAGAGTTCCTCGAAAACTATCTGCTGACTGATGAAGGTCTGGAAGCGGTTAATAAAGACAAAC
CGCTGGGTGCCGTAGCGCTGAAGTCTTACGAGGAAGAGTTGGCGAAAGATCCACGTATTGCCGC
CACTATGGAAAACGCCCAGAAAGGTGAAATCATGCCGAACATCCCGCAGATGTCCGCTTTCTGG
TATGCCGTGCGTACTGCGGTGATCAACGCCGCCAGCGGTCGTCAGACTGTCGATGAAGCCCTGA
AAGACGCGCAGACTGGCAGCGGCGGTAGCGGTGGCTCTCTGGAAGTGCTGTTTCAGGGCCCGAG
CGGTTCTGGTGGCAGCGGCAGCATTACCTATTTTTCGCTAGAATTTAGCATTCTGATGATCGCG
TTTTTTGCGATTTATTGGACCTTTAAAAACGATTATAAAATCCAGAACATTCTGATTCTGATTT
TCTCGTATATTATTTATATTCTGATAAACCCGTATTTCGCTCTGGTGCTGTTTATCTATACATT
TTTTATCCACTATTTCGCTTTACTAATTTTTGTACGGCGGAAGCGGTATATTTTCGCGACCTGT
ATGGCGTTTATTATTCTGAACCTGTGCTTTTTTAAATACTTTCCGTCCATAAAAGGCAGCGTGG
ATGAGATCCTGAACTTTTTCGGCCTGGAATTTCTGAACATTGATCTGGTTCTGCCAATCGGCAT
TAGCTTTTATACATTTACCTCGATAACCTATCTGGTAGAAGTGTATCAGAAACGGCGGCTGGAA
AGCTTTCTGAACCTGGCAACATTCCTGTCATTTTTTCCAACGCTACTGTCAGGACCGATTATGA
GGAGCTCGTTTTTCTTTGAACAAGCGTATCAGAAACGGGAATTTAAACATGCAAACCTGATAAT
AATATTACTGGTGTTTGGCATCGTGAAAAAAGTGCTGATCGCTAACTACCTGGGCATTTATGCG
AAAAGCATTCTGGACTTTCCACAGTCGTATAACTTCATCCAGCTGCTGTCCGCAATTTACGCTT
ATGCAATTCAGATCTACTGTGATTTTTCCGGCTATGTGGATCTGGTGTGCGCGTTTGCGCTGAT
GCTGGGCTTTACACTGCCCCCGAATTTTAATATGCCGTATCTGGCAAAAAATCTAAAAGATTTT
TGGGCGCGCTGGCATATAAGCCTGTCAACATTTATCCGCGATTATATATATATCCCGCTGGGAG
GGAACCGCAAAGGAATCCCACGGACCGTTGCGAACATCCTAATAGCGTTTATCCTGTCAGGCAT
GTGGCATGGGAACACGCTGGCGTTTATTGTGTGGGGCCTGTTACATGGCATTGGGATCGTGTTT
ATCCATTTACTAACCCTGTCCAAATTTAGCCTGCAGAAAATTCCAGCGCTGGGCCGCTTTCTGA
CATTTCAGTTTGTTTGCTTTACCTGGATTTTCTTTTATTATTCCAAAAACCTAGAAGATGCAAT
CGAATATTTTAAAGCGTGCTATTATAACTTCTTCCAGATTCCATCGTATAATGATATATATATG
TTAGTGGCGTTTGGAGTGCTGTTTATGATATATCCGCTGTTTATTAACTTTAAAGAATATTGTA
TTAAGATTCTGAACCTGACCCCATTTCTGCTAAAACCGTTTATAATCGCGTTTATTCTGCTGTT
AGTGTTTGCGTTTATGCCAGATGGCATTCCGGATTTTATTTATTCAAGCTTTTAATAA

**Fig. S14. DNA coding sequence for PelB-His$_{10}$-MBP-3C-*Cj*PatA expression construct.**

MKYLLPTAAAGLLLLAAQPAMAADSGSSGHHHHHHHHHHGSSGSKIEEGKLVIWINGD
KGYNGLAEVGKKFEKDTGIKVTVEHPDKLEEKFPQVAATGDGPDIIFWAHDRFGGYAQ
SGLLAEITPDKAFQDKLYPFTWDAVRYNGKLIAYPIAVEALSLIYNKDLLPNPPKTWEEIP
ALDKELKAKGKSALMFNLQEPYFTWPLIAADGGYAFKYENGKYDIKDVGVDNAGAKA
GLTFLVDLIKNKHMNADTDYSIAEAAFNKGETAMTINGPWAWSNIDTSKVNYGVTVLP
TFKGQPSKPFVGVLSAGINAASPNKELAKEFLENYLLTDEGLEAVNKDKPLGAVALKSY
EEELAKDPRIAATMENAQKGEIMPNIPQMSAFWYAVRTAVINAASGRQTVDEALKDAQ
TGSGGSGGSLEVLFQGPSGSGGSGSITYFSLEFSILMIAFFAIYWTFKNDYKIQNILILIFSYI
IYILINPYFALVLFIYTFFIHYFALLIFVRRKRYIFATCMAFIILNLCFFKYFPSIKGSVDEILN
FFGLEFLNIDLVLPIGISFYTFTSITYLVEVYQKRRLESFLNLATFLSFFPTLLSGPIMRSSFF
FEQAYQKREFKHANLIIILLVFGIVKKVLIANYLGIYAKSILDFPQSYNFIQLLSAIYAYAIQ
IYCDFSGYVDLVCAFALMLGFTLPPNFNMPYLAKNLKDFWARWHISLSTFIRDYIYIPLG
GNRKGIPRTVANILIAFILSGMWHGNTLAFIVWGLLHGIGIVFIHLLTLSKFSLQKIPALGR
FLTFQFVCFTWIFFYYSKNLEDAIEYFKACYYNFFQIPSYNDIYMLVAFGVLFMIYPLFIN
FKEYCIKILNLTPFLLKPFIIAFILLLVFAFMPDGIPDFIYSSF**

**Fig. S15. Amino acid sequence for PelB-His$_{10}$-MBP-3C-*Cj*PatA expression construct.**

### SI References


1. Chen, V. B., Arendall, W. B., Headd, J. J., Keedy, D. A., Immormino, R. M., Kapral, G. J., Murray, L. W., Richardson, J. S., and Richardson, D. C. (2010) MolProbity: All-atom structure validation for macromolecular crystallography. *Acta Crystallogr. Sect. D Biol. Crystallogr.* **66**, 12–21
2. Korlath, J.A., Osterholm, M.T., Judy, L.A., Forfang, J.C. and Robinson, R.A. (1985) A point-source outbreak of campylobacteriosis associated with consumption of raw milk. *J. Infect.Dis*. **152**, 592–596.
3. Miroux, B., and Walker, J. E. (1996) Over-production of proteins in *Escherichia coli*: Mutant hosts that allow synthesis of some membrane proteins and globular proteins at high levels. *J. Mol. Biol.* **280**, 289–298
4. Dempsey, J.A., Litaker, W., Madhure, A., Snodgrass, T.L., and Cannon, J.G. (1991) Physical map of the chromosome of *Neisseria gonorrhoeae* FA1090 with locations of genetic markers, including opa and pil genes. *J. Bacteriol.* **173**, 5476-5486.
5. Brott, A. S., Jones, C. S., and Clarke, A. J. (2019) Development of a high throughput screen for the identification of inhibitors of peptidoglycan *O*-acetyltransferases, new potential antibacterial targets. *Antibiotics*. **8**, 65
6. Claassens, N. J.; Siliakus, M. F.; Spaans, S. K.; Creutzberg, S. C. A.; Nijsse, B.; Schaap, P. J.; Quax, T. E. F.; van der Oost, J. (2017) Improving heterolgous membrane protein production in *Escherichia coli* by combining transcriptional tuning and codon usage algorithms. *PLoS ONE*. **12**(9), e0184355.
7. https://galaxyproject.org/use/codon-harmonizer/
